# Shaker it OFF: Biophysical Characterization of an Inactivating Potassium Conductance Mediating Object Segmentation in a Collision-Detecting Neuron

**DOI:** 10.1101/2025.09.04.674057

**Authors:** Gil Shaulsky, David Bellini, Dylan Ulloa, Eleni Nasiotis, Hala Khan, Hongxia Wang, Jiayi Luo, Herman Dierick, Chenghang Zong, Fabrizio Gabbiani, Richard B. Dewell

## Abstract

Homologous ion channels are expressed in sensory neurons across species where they shape responses to varied sensory stimuli, on a wide range of timescales. In grasshoppers, visual detection of looming stimuli which simulate an approaching predator involves the sub-cellular expression pattern of a newly characterized Shaker channel in a single identified neuron, discovered using a newly sequenced genome and single cell RNA sequencing. This channel shapes selectivity for approaching threats and was characterized with electrophysiological recordings, RNA interference, pharmacology and biophysical compartmental simulations. Our results explain how a slowly activating and inactivating potassium conductance common in neuronal dendrites contributes to visual object segmentation and implements a complex neural computation within a single neuron.

## Introduction

Animals must identify and discriminate external objects, such as an approaching predator, by segmenting them from their environment. This presumably requires discriminating between complex patterns of synaptic inputs on a wide range of spatiotemporal scales within dendrites of sensory neurons to generate an appropriate output and ultimately trigger behavioral responses. Synaptic integration depends on the differential expression of ion channels within a neuron’s dendritic arbor, which transform postsynaptic potentials as they propagate toward the spike initiation zone. Across species, neurons use evolutionarily conserved ion channel genes (Jegla et al. 2009), whose differential expression sets up many types of information-shaping processes. These genes are involved in processing diverse sensory stimuli such as spectro-temporally modulated sounds or dynamic visual patterns, even though they originated in organisms without ears or eyes (Salkoff et al. 1992; Jacobs et al. 2007; Jegla et al. 2009).

Therefore, insights into the role of channels in one organism likely have implications across distantly related species.

Voltage-gated ion channels dynamically change their conductance based on time-varying membrane potentials and differentially affect synaptic inputs depending on their time course and location. Some of these channels inactivate, which prevents current from flowing. Inactivation may be voltage-dependent and is usually slower than activation (Jegla et al. 2009). This difference in timing results in current passage for a limited time upon depolarization, as in the fast sodium channels that generate the rising phase of an action potential (Hodgkin and Huxley 1952).

The *Shaker* gene family defines a set of voltage-gated potassium channels, some of which inactivate. These genes were first characterized in fruit flies (*Drosophila melanogaster*), but homologous ones are present in many species, including the mammalian Kv1 subfamily (Jan et al. 1977; Curran et al. 1992). In flies, one role of Shaker channels includes regulating the time course of contrast-specific visual responses (Gür et al. 2020). Fast-inactivating potassium channels have clearly defined roles in neuronal computations such as spike frequency adaptation and f-I curve smoothing (Hoffman et al. 1997). Less is known about the role of slowly-inactivating dendritic potassium channels besides their involvement in delaying spiking (Storm 1988).

Dendritic potassium channels have been implicated in selectivity of predator escape behavior (Dewell and Gabbiani 2018). The American bird grasshopper *Schistocerca americana* reacts to a looming stimulus simulating an approaching predator by jumping and flying away (Fotowat and Gabbiani 2007). There is selective pressure to reliably escape imminent predation, but not to jump at other times since this would break the grasshopper’s camouflage (Mitra et. al. 2025). Indeed, these animals consistently jump in response to a looming stimulus, but not other stimuli with features in common, such as those approximating birds flying nearby or clouds passing in front of the sun (small moving edges or global luminance changes, respectively).

A large, identified neuron in the grasshopper optic lobe, the Lobula Giant Movement Detector neuron (LGMD; O’Shea and Williams 1974), processes looming stimuli and drives escape behavior (Fotowat et al. 2011). It lies three synapses away from the photoreceptors and receives input from every facet in the eye (Figure 1A; Krapp and Gabbiani 2005). In response to a looming stimulus, the LGMD generates a gradual increase in firing followed by a sharp drop before the moment of escape (Fotowat et al. 2011). The LGMD projects to a dedicated output neuron, the Descending Contralateral Movement Detector (DCMD), with which it has 1:1 spike parity (O’Shea and Williams 1974). The DCMD in turn projects to flexor and extensor motor neurons of the hind legs that cause muscle co-contraction and then release of the legs when firing drops, allowing the animal to jump away (Burrows and Rowell 1973; Fotowat and Gabbiani 2011).

**Figure 1.**
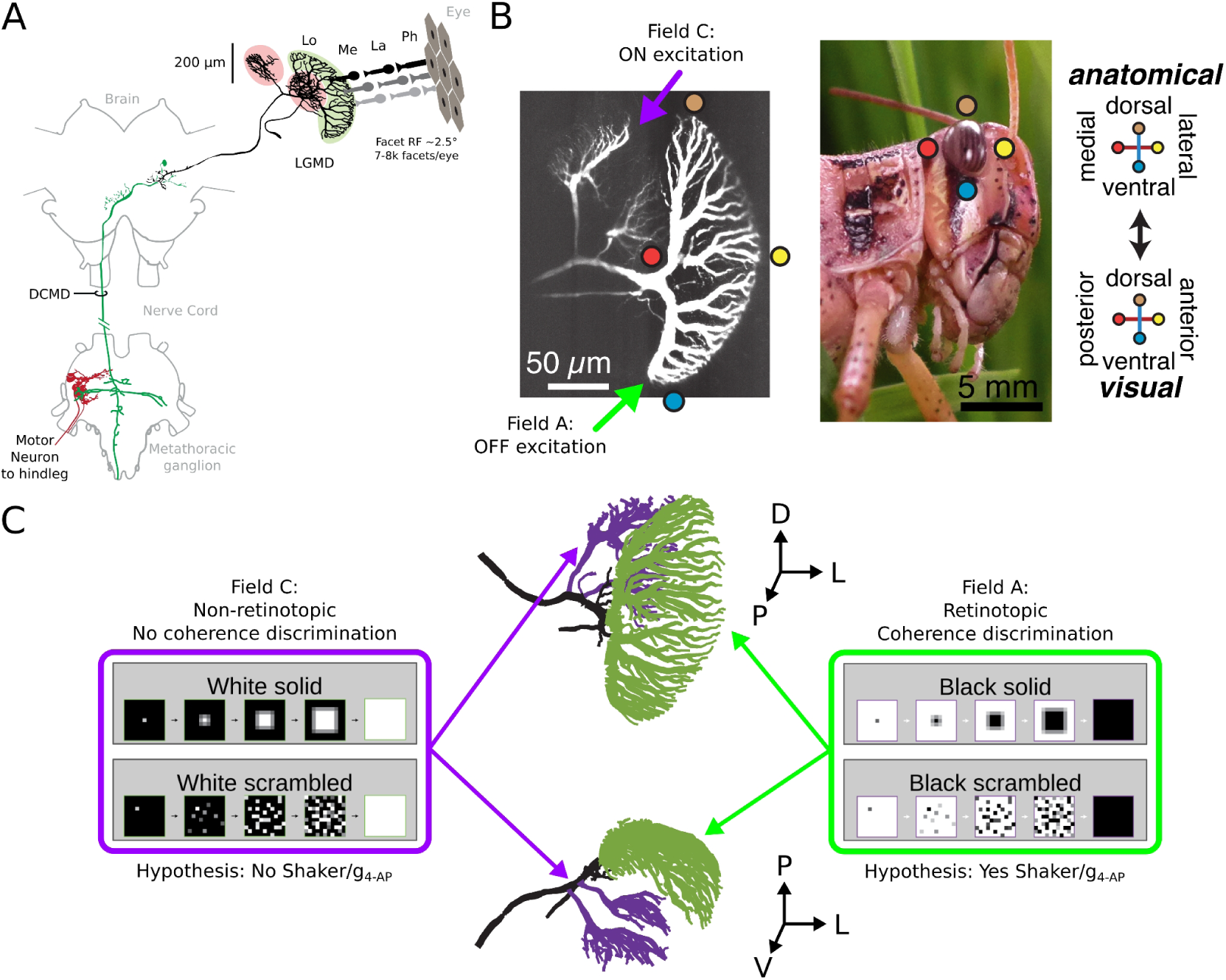
Coherence discrimination in the LGMD is contrast polarity dependent due to segregation of inputs, coinciding with subcellular patterns of channel expression. A) Diagram of the grasshopper visuomotor pathway for escape jumps. The adult compound eye is composed of 7-8 thousand individual facets with individual receptive fields (RF) of about 2.5°. Ph – photoreceptor layer, La – lamina, Me – medulla, Lo – lobula layers of the optic lobe. In each lobula, the LGMD receives input from every facet and projects to a dedicated output neuron in the brain, the Descending Contralateral Movement Detector (DCMD). In the metathoracic ganglion, the DCMD innervates hindleg motor neurons that produce jumps. B) Left: Maximum projection of a 2-photon stack of a fluorescent stained LGMD. The largest part of the dendrite, field A, receives OFF (black) input from every facet of the compound eye in a retinotopic pattern (colored circles). ON (white) input from every facet goes to the smaller field C without retinotopic organization. Right: Side view *Schistocerca americana* with diagram of the retinotopic mapping of the Lobula Giant Movement Detector Neuron (LGMD). C) Hypothesis for the localization of the 4-AP-sensitive conductance contributing to contrast and coherence detection in the LGMD. Based on earlier work (Dewell et al. 2022), we hypothesized that a 4-AP-sensitive K^+^ conductance is localized in Field A since it receives retinotopic OFF inputs and implements object segmentation. Conversely, we hypothesize that Field C lacks the 4-AP-sensitive conductance since it receives non-retinotopic ON inputs and does not implement object segmentation. D – dorsal, P – posterior, V – ventral, L – lateral. A and B are adapted from Dewell and Gabbiani 2018. C is adapted from Jones and Gabbiani 2010.

The dendritic arbor of the LGMD is divided into distinct fields. The largest, field A, receives OFF (black) retinotopic excitation from every facet (Figure 1B; Peron et al. 2009; Zhu and Gabbiani 2016). In contrast, ON (white) excitation goes to the smaller field C, without retinotopic organization (Dewell et al. 2022). The retinotopically-organized inputs to the LGMD’s dendrite are shaped by the various conductances expressed therein, and that interaction determines the cell’s spiking output. A hyperpolarization-activated, cyclic-nucleotide-gated (HCN) cation conductance selectively potentiates responses to spatially coherent black visual stimuli (Dewell and Gabbiani 2018). Conversely, an inactivating voltage-gated potassium conductance blocked by 4-aminopyridine (4-AP) decreases the response to black stimuli, with the greatest reduction for spatially incoherent ones consisting of the same elementary luminance changes as looming stimuli. The 4-AP-sensitive conductance (g_4-AP_) contributes to looming stimulus segmentation through spatial coherence detection (Dewell and Gabbiani 2018). However, the molecular identity, kinetics, and the dendritic localization of g_4-AP_ remain unknown (Figure 1C).

We used electrophysiology, pharmacology, single-cell LGMD RNA sequencing, genetic manipulation and biophysical modeling to identify g_4-AP_ as a newly-discovered Shaker channel and to characterize its role in spatial coherence selectivity for looming stimuli. We showed that the resulting conductance is only present in field A, and that its voltage dependence and time constant for activation and inactivation are within the behaviorally relevant range for approaching predator detection. Furthermore, we demonstrated that this channel plays a role in spike frequency adaptation and that its implementation in an existing LGMD biophysical model contributes to approaching object segmentation. Similar ion channels could potentially implement related neural computations across the animal kingdom.

## Methods

### Animals

American bird grasshoppers (*Schistocerca americana)* were reared as previously described (Dewell and Gabbiani 2018). For visual response and electrophysiology experiments, female animals ∼10 days after final molt were selected, excluding those with obvious eye defects or sluggish movement in their home cage. Animals in behavioral experiments had the additional criterion of no missing or deformed legs. For RNA interference (RNAi) experiments, female animals within the first week after final molt were used due to the required incubation time.

### Dissection

Animals were prepared as previously described (Dewell and Gabbiani 2018). Legs, wings, antennae, and jaws were removed, then the animals were placed in a 3D-printed plastic stage allowing to bathe the head in locust saline (140 mM NaCl, 5 mM KCl, 5 mM CaCl_2_, 5 mM MgCl_2_, 4 mM NaHCO_3_, 6.3 mM HEPES, 73 mM sucrose, pH 7.0). The abdomen was not submerged, allowing the animal to breathe. An incision was made in the neck membrane, through which the gut was removed. The head was tilted forward 90° and secured to the side of the stage with wax. Muscle, fat, and air sacs were removed posterior to the right optic lobe, preserving tracheas going to the lobe and retina. Finally, the protective brain sheath around the lobe was removed mechanically, exposing the neural tissue.

### DCMD Recording

Two wires (stainless steel 304, Formvar insulation, 420 Ω*cm/ft, California Fine Wire cat. #100192) were bent into hooks, with a small deinsulated area on the inside of each crook. These hook electrodes were placed around the left nerve cord (contralateral to the right eye and optic lobe), with the deinsulated part on the medial side, slightly dorsal to match the location of the DCMD within the nerve cord. The difference in voltage between the two hooks was amplified relative to a ground electrode and filtered by an A&M Systems differential AC amplifier (Model 1700), then further amplified by a Brownlee Precision instrumentation amplifier (Model 440), and digitized and stored on a computer by a PowerDaq data acquisition card (PD2-MF-16-500/16H). The position of the hooks was adjusted until the DCMD spikes had the highest amplitude and were clearly distinguishable from other nerve cord spikes.

### LGMD Staining

Borosilicate thin-walled glass pipettes with filaments (1.2 mm OD, 0.9 mm ID, World Precision Instruments, TW120F-4) were prepared on a micropipette puller (Sutter Instrument Co. Model P-97), resulting in a thin tip with a ∼1 cm long taper. For staining, the tip was filled with 0.4-1.2 µL of 10 mM fluorescent dye (Alexa 594 hydrazide salt, Sigma or Atto 594 hydrazide salt, Atto-Tech), then backfilled with a mix of potassium chloride (1.8 M) and potassium acetate (1.2 M). The two solutions in the tip were allowed to partially equilibrate over the course of 5-30 min. Electrodes were placed into a holder with a silver wire coated in a thin layer of silver chloride (PPH-1P-BNC; ALA Scientific Instruments). A reference wire of the same composition was placed in the bath. Signals from the electrode were amplified by an intracellular amplifier (AxoClamp-2A or NPI SEC-10LX), then by a Brownlee amplifier (Model 440). The tip of the electrode was lowered into the lobula of the exposed right optic lobe until it entered the LGMD, indicated by 1:1 spike parity with the extracellular DCMD recording.

Negative current pulses were applied to inject the electronegative dye molecule into the cell by iontophoresis. Fluorescence was excited with 543-578 nm light from a Leica EL6000 external light source. Epifluorescence light was passed through a Texas Red filter cube and imaged with an iDS digital camera (UI-3240CP-M). Dye was injected only until the entire LGMD dendritic arbor was dimly visible (∼30 seconds cumulative with current on). Following withdrawal of the staining electrode from the LGMD, the stain was used to guide subcellular targeting for subsequent recordings or drug application.

### LGMD Targeted Recordings

After staining as described above, another borosilicate pipette was pulled as above, then filled with a potassium chloride (3 M) solution pre-mixed with a small amount of fluorescent dye (Alexa or Atto, ∼0.2-1% by volume), with a resistance of 10-15 MΩ. This recording electrode was attached to the holder and connected to amplifiers as described above, placed above the desired area of the LGMD, and lowered until it broke into the cell, as described above. For input resistance characterization and evoked firing experiments, the recording amplifier was placed into Discontinuous Current Clamp mode (DCC), positive and negative current pulses applied, and the resulting spiking and voltage response recorded. For ion channel kinetics characterization, the recording amplifier was placed in Single Electrode Voltage Clamp mode (SEVC), gradually increasing clamp gain as high as possible without causing stereotypical high-amplitude, high-frequency oscillations (minimum gain of 5 µA/V). For activation kinetics characterization, the cell was clamped at a single hyperpolarized pre-step potential, followed by a depolarizing step of varied magnitude while the current response was recorded. For inactivation kinetics characterization, the cell was clamped at a varied hyperpolarized pre-step potential followed by depolarizing steps to the same potential. While completely space-clamping the cell would be impossible, we estimated the average steady-state voltage change relative to resting membrane potential to be 85-93% across dendritic field A using the methods described in Dewell and Gabbiani (2018).

### Visual Stimuli

Looming and translating visual stimuli were programmed in C or Matlab with Psychtoolbox-3 and displayed on an LG StudioWorks 221U, 21-inch CRT (200 frame/s) or ASUS ROG Strix XG249CM LCD monitor (240 frame/s). The animal was positioned 180 mm from the screen and aligned to its center. Looming stimuli had an *l*/|*v*| value of 50 ms (Gabbiani et al., 1999). Coarse black looms, pixelated at the size of photoreceptor’s receptive fields were used (Jones and Gabbiani 2010). The coherence of coarse looming stimuli was varied from fully coherent (100%) to fully random (0%) with intermediate values of 80%, and 60% coherence by adding zero-mean spatial jitter to the coarse pixel locations (Dewell and Gabbiani, 2018). We also used coarse white looms at two coherences (100%, 0%), and translating stimuli (bars with a long axis spanning the screen and a short axis spanning ∼5 degrees in visual space (16 mm on the screen). For visual response experiments, stimuli were separated by a 2-minute recovery period, during which the animals were presented with other sensory stimuli to reduce the effect of habituation (turning the lights on and off, jingling keys, and/or gently stroking the abdomen).

### Pharmacology

4-aminopyridine (4-AP) was purchased from Sigma Aldrich. It was dissolved in distilled water to a concentration of 200 µM. For puff application, it was further diluted to 10 µM in locust saline containing a small amount of Fast Green FCF dye (1.24 µM), and then loaded into a borosilicate pipette with a ∼2 µm opening. This drug pipette was attached to an air source and a pressure regulator (World Precision Instruments, PV830, Sarasota, FL). It was then lowered until the tip was just inside the targeted neural tissue, guided by the fluorescent LGMD stain (see above for staining procedure). Using the minimum required pressure, drug solution was injected into neural tissue proximal to the appropriate part of the dendrite, with the Fast Green indicating how far the drug spread. For iontophoresis, 4-AP at 200 µM in distilled water only was loaded into a similar pipette with a ∼1 µm opening, which was then placed in an electrode holder attached to an AxoClamp amplifier (as above). The pipette was lowered into the tissue, guided by the fluorescent stain, then a high current (∼100 nA) was applied for 15 seconds. The same pipette was subsequently used to apply 4-AP to field A as a within-animal positive control. For bath application, 4-AP was diluted in locust saline to a larger volume (150-1000 µL) for better mixing, then added to the bath via a syringe for a final concentration of 571 µM with 1.43 mM mecamylamine for voltage-clamp experiments or 171 µM alone for some visual experiments. Mecamylamine was used to block visual inputs, as the 4-AP-mediated increase in synaptic inputs made it harder to maintain a stable voltage-clamp recording. The same amount of bath solution was subsequently removed to keep total saline level constant.

### Single-cell Sequencing and Transcript Mapping

For single-cell mRNA sequencing of the LGMD, animals were prepared and the LGMD was injected with fluorescent dye as described above until the cell body was visible. Tissue above (posterior to) the cell body was removed with forceps, and a glass pipette was placed next to it. Using gentle negative pressure to prevent completely aspirating the cell body, it was adhered to the opening of the pipette. The pipette and cell body were withdrawn together, causing the body to separate from the rest of the cell. The tip of the pipette was placed in a buffer solution and gentle positive pressure was applied to release the cell body. The cell bodies were then processed using snapTotal-seq as described in Niu et al., 2024.

After sequencing, data was mapped to the *Schistocerca americana* genome using the STAR aligner (Dobin et al. 2013) and the count matrix generated using featureCounts (Liao et al. 2014). Then, counts per million were counted using the EdgeR package in R (Robinson et al. 2010). The resulting values were used to identify the gene most likely responsible for the 4-AP-sensitive conductance: a Shaker gene on chromosome 4. See Supplemental Table 1 for a list of relevant potassium channel genes and their expression level in *S. americana*.

### Phylogeny

Homologous proteins from common model species across multiple phyla were found by using the BLASTP program and the ClusteredNR database (NCBI; Altschul et al. 1997). For each species, the protein with highest percent identity to the corresponding protein in *S. americana* was selected and transmembrane regions identified by DeepTMHMM (Hallgren et al. 2022). The regions of those sequences between the first and last transmembrane helices were used to generate a phylogenetic tree, ordered by the Fast Minimum Evolution method, with branch length determined by the Grishin method (Grishin 1995; Desper and Gascuel 2004). Graphical representation of the tree was performed using IcyTree (Vaughan 2017). See Supplemental Table 2 for the regions used.

### RNA Interference

Following the recommendations in the MEGAscript® RNAi Kit (Invitrogen), a double-stranded RNA (dsRNA) probe targeting a 580 bp region common to all known transcripts of the *Shaker* gene in *S. americana* was generated. To enable *in vitro* transcription, a T7 RNA polymerase promoter sequence (5’-TAATACGACTCACTATAGGGAGA-3’) was appended to the 5’ end of opposing gene-specific primers (Forward: 5’-CGAGTACTTCTTCGACCGC-3’; Reverse: 5’-CGACTGTCGCTAGGGTGATG-3’, see Supplemental Figure 1). PCR amplification was performed using the 2x Rapid Taq Master Mix (Vazyme) according to the manufacturer’s instructions, with the following modifications: 0.5 µg of cDNA template, an annealing temperature of 63 °C, a 5-second extension time, and 35 cycles. The resulting amplicon was visualized on a 1.5% agarose gel, confirming the expected product size. The remaining PCR product was purified using the QIAquick PCR Purification Kit (Qiagen) and eluted with 30 µL of pre-warmed (50 °C) nuclease-free water following a 10-minute incubation. For transcription, 0.4 µg of purified DNA was used in a T7 polymerase reaction incubated at 37 °C for 4 hours. The resulting RNA product was treated with nucleases and then purified according to the MEGAscript protocol. The purified RNA was eluted with two separate 75 µL volumes of preheated (95 °C) elution buffer. The dsRNA was diluted to 12 ng/µL using locust saline. 180 ng (15 µL) total dsRNA was injected into healthy adult females within a week of their final molt, using a 22s gauge curved, beveled needle placed between the ventrolateral thorax and first abdominal segment, avoiding the gut and nerve cords. Experimental animals were group-housed and given two additional boosters on the 3^rd^ and 6^th^ days after the first injection, using the same volume and protocol. Recordings were done on the 9^th^ day after initial injection. After an animal’s recordings were complete, the brain, optic lobes, and retinas were removed and frozen in liquid nitrogen for RNA analysis.

### Quantitative PCR (qPCR)

Neural tissue was stored at-80 °C. Total RNA was extracted using a rotor-stator homogenizer in combination with the PureLink® RNA Mini Kit (Invitrogen).

Residual genomic DNA was removed by DNase treatment using the TURBO DNA-free™ Kit (Invitrogen). First-strand cDNA was synthesized from 50 ng of total RNA using random hexamer primers and the SuperScript™ IV First-Strand Synthesis System Kit (Invitrogen). Quantitative real-time PCR (qPCR) was performed using iTaq™ Universal SYBR Green Supermix (Bio-Rad) with gene-specific primers to assess RNAi knockdown efficiency. *Shaker*-specific primers (Forward: 5’-GGCATTCTGCTAACGAGGG-3’; Reverse: 5’-GTCTCAATGCTCATCGCG-3’, see Supplemental Figure 1) and GAPDH-specific primers (Forward: 5’-TCCAAGTGCAGATGCCCCAA-3’; Reverse: 5’-CTTTTGCCAGTGGTGCCAGG-3’) Annealing temperature for qPCR: 58 °C.

### Data Analysis

Data from visual and physiological experiments were analyzed with custom Matlab code (MathWorks, Natick, MA). For responses to visual stimuli, DCMD spikes were identified via a spike sorting algorithm based on a manually set threshold. The raster of these spikes was convolved with a zero-mean Gaussian with standard deviation of 20 ms to yield the instantaneous firing rate (IFR) at each time point during the stimulus (Gabbiani et al. 1999). For voltage and current clamp data, response traces from individual trials were filtered through a one-dimensional median filter to remove spikes (symmetric window size: 24.909 ms, sample rate: 40.146 sample/ms). All traces for each stimulus were time-aligned to the start of the command step and, for all trials except inactivation characterization, were also zeroed to the baseline voltage or current before the start of the step. The average response value at each time point was calculated for each drug condition to give the mean control and drug response time courses. Subtracting the drug trace from the control one yielded the mean difference trace, representing the 4-AP-sensitive component of the control response. Much of the data was non-normal, so non-parametric tests were used. For paired, within-animal comparisons, such as responses before and after drug application, a Wilcoxon Sign Rank (WSR) test was used. For unpaired data, such as comparing RNAi-treated animals to wild-type, a Wilcoxon Rank Sum (WRS) test was used.

To isolate the effect of voltage on channel gating only, the conductance of the population of 4-AP-sensitive channels (in units of mS) was calculated as g_4-AP_ = I_4-AP_/(V_step_-E_K_), where I_4-AP_ is the mean peak 4-AP-sensitive current across animals in nA, V_step_ is the voltage of the depolarizing step in mV, and E_K_ is the reversal potential of potassium in the LGMD. Under the assumption that the lowest calculated conductance represented zero channels active and the highest represented all channels active, we calculated percent activation from the conductance at each depolarizing step by subtracting the lowest, then dividing the result by the highest. The same voltage dependence values were calculated for inactivation, using the constant V_step_ =-35 mV when calculating conductance. When plotting voltage (step or pre-step) against the corresponding activation percentage, the resulting data are roughly sigmoid, typical for activation of voltage-gated channels (Storm 1988).

### Fitting

To estimate percent activation at other voltages, these data were fitted with the least-squared-error method (Matlab), which was then used to find the half maximum voltage for activation and inactivation and the percent permissive at rest.

Since the time constants for both activation and inactivation gating depend only on the step (and not pre-step) voltage, the currents from the activation voltage dependence characterization experiments (variable step, constant pre-step) were sufficient. A double exponential of the form:

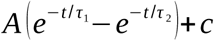

was used for fitting, where t is time after the start of the voltage step, c is the baseline current, and 𝜏_1_ and 𝜏_2_ are the time constants of the falling and rising phases of the 4-AP-sensitive current traces for each step. The fit procedure was applied to each animal’s mean 4-AP-sensitive current traces at depolarizing steps between-50 and-20 mV. These steps were the only ones where the 4-AP-sensitive inactivating current in most animals was large enough to fit. Only calculated time constants between 4 milliseconds and 10 seconds were included.

### Simulations

All simulations were performed in NEURON (Yale), using the LGMD model available on ModelDB (model # 2019879).

### Notation

Standard deviations are abbreviated by s.d. and the standard errors of the mean by s.e.m.

## Results

The analysis presented here characterizes the genetic identity, dendritic localization, and kinetics required to explain the role of a voltage-gated channel in a single-neuron computation part of a pathway critical to escape behavior (Figure 1A). In earlier work, simulations found that addition of a slowly-inactivating voltage-gated potassium channel to a multi-compartment NEURON model of the LGMD was necessary to obtain black visual looming stimulus responses sensitive to spatial coherence (Dewell and Gabbiani 2018). In addition, 4-aminopyridine (4-AP), a blocker of inactivating voltage-gated potassium channels, has been shown to differentially affect responses to black visual stimuli, selectively potentiating LGMD responses to spatially-incoherent looming patterns (Dewell and Gabbiani 2018). Those inputs are mapped retinotopically onto a dedicated dendritic subfield of the LGMD (Figure 1B). In contrast, white looming visual stimuli that are processed in a distinct dendritic subfield lacking retinotopy, generate responses insensitive to stimulus coherence (Dewell et al. 2022). Taken together, these results suggest the presence of an uncharacterized 4-AP-sensitive, slowly-inactivating potassium conductance g_4-AP_ in dendritic subfield A of the LGMD that is active during looming stimuli, whereas field C is expected to lack that conductance (Figure 1C).

### A Shaker channel is expressed in the LGMD

To find the channel gene(s) underlying this 4-AP sensitive conductance, we sequenced the mRNA found in five LGMD cell bodies and compared the results with a recently published high-quality *Schistocerca americana* genome to identify candidate channel genes. Multiple voltage-gated potassium channel genes were discovered, including ones from the *Shaker* and *Ether-a-go-go* families (see Supplemental Table 1). Those channel genes had high sequence homology with their counterparts in other animals, including non-arthropods, to the point where *Shaker* sub-family genes (*Shaker*, *Shab*, *Shal*, and *Shaw*) cluster with themselves, rather than with their organism, when using the Fast Minimum Evolution method (Figure 2A). The K^+^ channel gene that was most highly expressed in the LGMD was located on chromosome 4 and coded for a Shaker channel (homologous to Kv1 in mammals). The channel sequence included six residues near the cytoplasmic ends of the S5 and S6 transmembrane helices that are necessary for 4-AP binding in *Drosophila* Shaker channels (Figure 2B; Pinto-Anwandter 2024).

**Figure 2.**
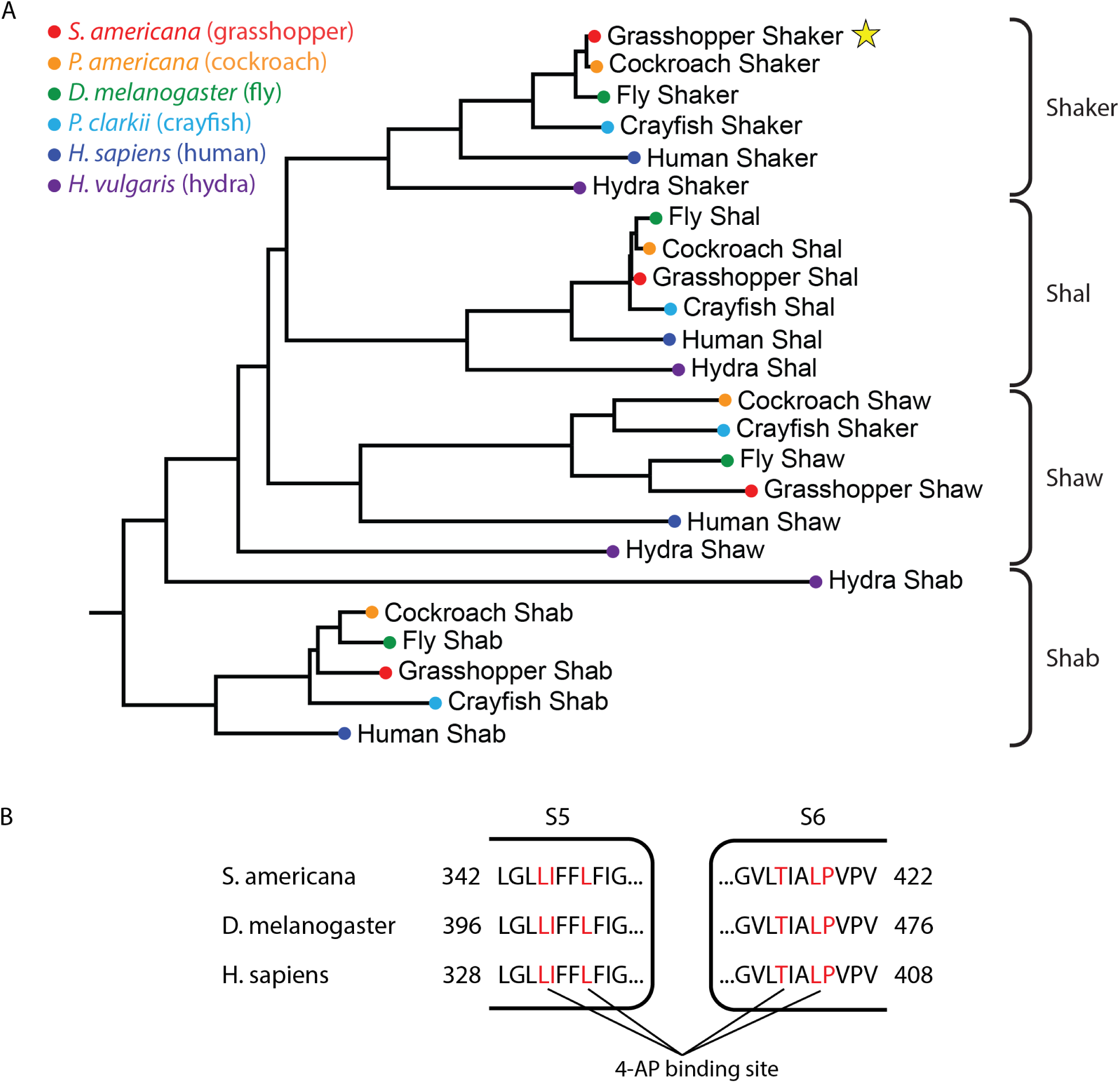
A novel grasshopper Shaker channel with a canonical 4-AP binding site is expressed in the LGMD. A) Phylogram of four Shaker-family potassium channels (Shaker, Shab, Shal, and Shaw) across six animal species, generated using the Fast Minimum Evolution method. Branch length is proportional to relative evolutionary distance, determined by fraction of mismatched amino acids (Grishin model). Species are grouped by color, with braces indicating clustering by channel type. Star indicates *S. americana* Shaker. B) 4-AP binding location in *S. americana, D. melanogaster*, and *Homo sapiens* Shaker/Kv1 show conserved binding sequence. In models of the *D. melanogaster* Shaker channel, the cytoplasmic ends of the S5 and S6 transmembrane helices are close together and the residues labeled in red form a hydrophobic pocket where 4-AP binds.

We also found two accessory protein genes in the LGMD that may affect the resulting conductance: a Shaker β subunit and a Discs-Large Homolog (DLG) gene. Some Shaker/Kv1 channels inactivate only in the presence of a β subunit, while the DLG gene encodes a membrane-associated guanylate kinase whose mammalian homolog PSD-93 regulates the subcellular localization of voltage-gated potassium channels (Rettig et al. 1994; Morales et al. 1995; Ogawa et al. 2008).

### Blocking voltage-gated potassium channels only in dendritic field A of the LGMD selectively potentiates incoherent black loom responses

In *S. americana*, the magnitude of the LGMD response and the probability of evoking a jump to a black looming stimulus increase with spatial coherence (Dewell and Gabbiani 2018). As stated above, the application of 4-AP reduces spatial selectivity by increasing responses to black incoherent stimuli. Since excitatory synaptic inputs originating from OFF (luminance decrement) stimuli project onto field A in a retinotopic pattern, but ON-sensitive inputs project to field C without retinotopy, we predicted that 4-AP-sensitive potassium channels are expressed in field A only, where they selectively attenuate responses to black incoherent stimuli. DCMD firing in response to looming stimuli was recorded and used to estimate the instantaneous firing rate (IFR) at each time point (Figure 3A-D).

**Figure 3.**
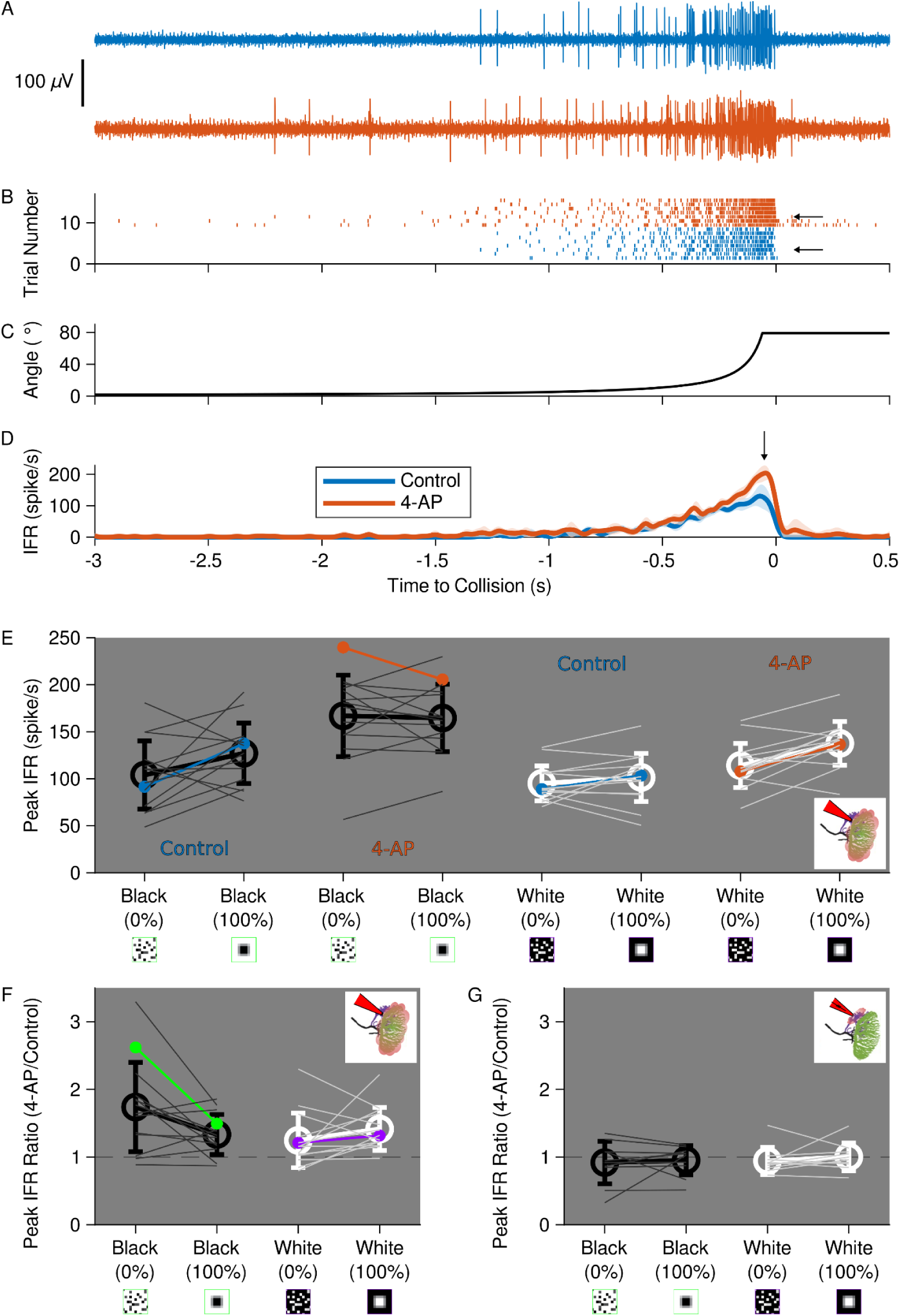
Expression of a 4-AP-sensitive conductance in field A only is necessary for coherence discrimination. A) Example DCMD responses to a solid black loom before (top, blue) and after (bottom, red) 4-AP application. B) Rasters of DCMD spikes to each trial of the same animal as (A) colored as above. Black arrows indicate the trials in A. C) Angular size time course for a solid loom with an *l*/|*v*| of 50 ms, maximum ∼80°. D) Mean ± s.d. of the instantaneous firing rate (IFR) of responses shown in (B). E) Effect of stimulus coherence (0% or 100%) on peak IFR, before and after addition of 4-AP to field A. Thin lines are individual animals, colored lines are the examples used in A-D (blue: control, orange: after 4-AP), and thick lines represent mean across animals. All stimuli had significant increases after application of 4-AP (see Statistics Table). F) Effect of stimulus coherence on the ratio of peak IFR for each stimulus before and after application of 4-AP to field A. Thin lines are individual animals, with colored lines for the example used in A-D. Thick lines are mean across animals ± s.d. Asterisk indicates a significant difference (black: p = 0.0419, white: p = 0.1726, WSR). Dotted line represents a ratio of 1 (i.e., no change). G) Effect of stimulus coherence on the ratio of peak IFR for each stimulus before and after application of 4-AP to field C. Thin lines are individual animals; thick lines are mean across animals ± s.d. No significant difference was observed (black: p = 1, white: p = 0.375, WSR).

After 4-AP application to field A, all stimuli had a significantly higher peak firing rate, but the net increase was the greatest for black incoherent stimuli (Figure 3E). The ratio of peak instantaneous firing rate (IFR) after versus before 4-AP application was significantly higher for black stimuli with 0% coherence than for those with 100% coherence (Figure 3F, p = 0.042, WSR). While responses to both 0% and 100% coherent white stimuli exhibited increased firing, the relative increase was about the same, (Figure 3F, p = 0.17, WSR). This suggests that while the 4-AP sensitive conductance reduces activity in response to all visual stimuli, it selectively weakens responses to black incoherent stimuli relative to coherent ones.

The increase in responses to all stimuli is likely due to an increase in input resistance, from 3.8 ± 0.1 MΩ to 5.5 ± 0.1 MΩ (mean ± s.d., n = 6, p = 0.031, WSR, Supplemental Figure 2). This is caused by a reduction in the membrane’s active channels and means that the same excitatory postsynaptic currents (EPSCs) result in larger excitatory postsynaptic potentials (EPSPs). Since the potassium reversal potential in the LGMD is below the resting membrane potential, as it is in most neurons, 4-AP raised the LGMD resting membrane potential from-64.7 ± 2.1 mV to-62.4 ± 3.6 mV (mean ± s.d., n = 6, p = 0.094, WSR, Supplemental Figure 3). This puts the LGMD closer to its spike threshold, so the same size EPSPs elicit more firing. These changes are presumably sufficient to produce a general strengthening of visual responses, suggesting that the 4-AP-sensitive conductance has a suppressive effect, scaling down the LGMD’s visual responses. However, an increase in input resistance and resting membrane potential do not explain the greater relative increase in responses to incoherent black stimuli, which corresponds to the greater suppression of those responses by the 4-AP-sensitive conductance in the control condition. Since by design the difference must be based on the way the LGMD combines the spatiotemporal patterns of EPSPs resulting from coherent versus incoherent black stimuli, elucidating this mechanism required characterizing the kinetics of the conductance (see below).

In contrast, when 4-AP was applied to field C, there was no significant increase in responses for any of the visual stimuli tested, indicating that the 4-AP-sensitive conductance is not active there (Figure 3G). These results are consistent with the presence of a 4-AP-sensitive potassium conductance that is functional in field A, where it plays a specific role in coherence selectivity for black stimuli, and that is not at all functional in field C.

### RNAi knock-down of *Shaker* changes LGMD coherence preference and decreases 4-AP sensitivity

Based on the pharmacology and gene expression data, the 4-AP sensitive conductance localized in field A is likely produced by Shaker channels. We tested the role of the *Shaker* gene in visual responses with RNA interference (RNAi) to knock down its expression. After dsRNA injection and 10 days of incubation to allow for channel turnover, animals were dissected and prepared as above with the LGMDs stained and DCMD responses to visual stimuli recorded before and after 4-AP application (Figure 4A-D). At the end of recording, neural tissue was extracted and Shaker mRNA level was quantified. Since blocking the 4-AP-sensitive conductance did not have an effect on coherence selectivity for white stimuli, only black stimuli were tested. Stimuli included 100% and 0% coherent black looms as well as translating black bars on a white background, since earlier experiments suggested that 4-AP also potentiates LGMD responses to translating stimuli (Dewell and Gabbiani 2018). Bars moving in the four cardinal directions were presented in a random order, interspersed with looming stimuli. All translation directions produced similar responses (initially high firing that quickly attenuated to almost none), but since the stimulus monitor was wider than it was tall, left and right translations occurred over a longer period of time than did up and down translations, so vertical and horizontal responses were binned separately.

**Figure 4.**
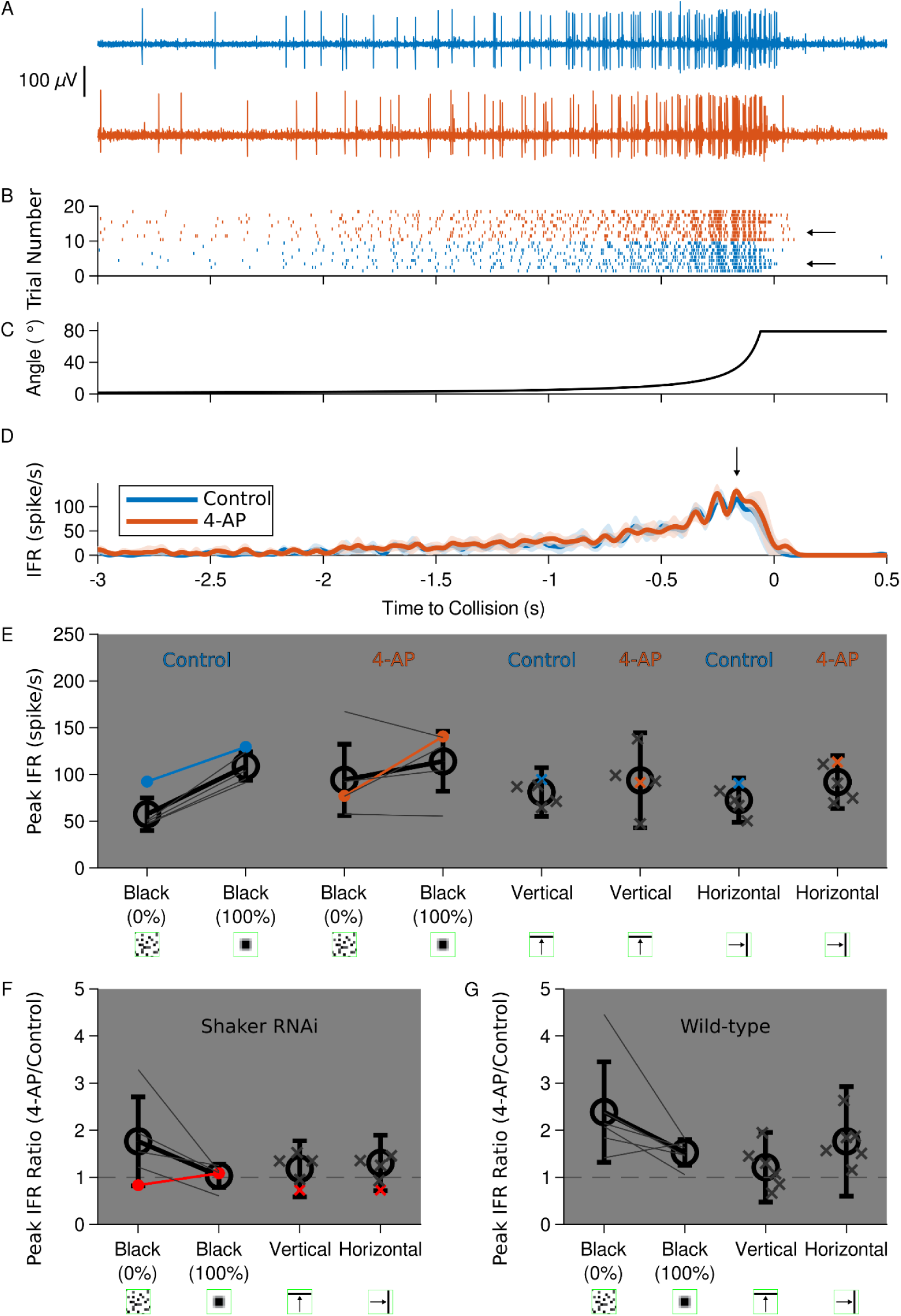
RNAi of Shaker changes visual response properties and reduces 4-AP sensitivity. A) Example DCMD response to a solid black loom in a Shaker knock-down animal. Top: control, bottom: after 4-AP application. B) Rasters of DCMD spikes to a solid black loom for one Shaker knock-down animal. Blue: control, orange: after 4-AP application to field A. Black arrows indicate the trials in A. C) Angular size time course for a solid loom with an *l*/|*v*| of 50 ms, maximum ∼80°. D) Mean IFR ± s.d. to a solid black loom for one Shaker RNAi animal. Blue: control, orange: after 4-AP application to field A. Black arrow indicates the time of peak firing. E) Effect of stimulus coherence (0% or 100%) and translating stimulus direction (Vertical, Horizontal) on peak IFR, before and after addition of 4-AP to field A of a Shaker knock-down animal. Thin lines and solid marks are individual means, colored lines and dots are the examples used in A and B (blue: control, orange: after 4-AP), and thick lines and open circles are population means. F) Effect of stimulus coherence on the ratios of peak IFR before and after application of 4-AP to field A of a Shaker knock-down animal, plotted as above. Red indicates the example animal used in 4A-D. Dotted line represents a ratio of 1 (i.e., no change). G) Effect of stimulus coherence on the ratios of peak IFR before and after application of 4-AP to field A of a wild-type animal, plotted as above.

Expression levels in RNAi-treated animals were reduced to 24% ± 4% (mean ± s.d., n = 5, see Supplemental Figure 4), of the average level in wild-type. Compared to those wild-type animals, whose LGMDs had a near-zero resting firing rate, *Shaker* RNAi-treated animals exhibited higher spontaneous LGMD firing when no visual stimulus was being presented (though not significantly, p = 0.33, WRS; n = 5 wild type, n = 5 RNAi; see Supplemental Figure 5). 4-AP increases spontaneous firing in both cases, but the difference is smaller for the RNAi animals (though not significantly, p = 0.095, WRS). They also had higher peak firing rates in response to translating stimuli (though not significantly, p = 0.082, WRS, Figure 4E). In contrast, responses to looms were slightly lower than in wild-type animals, possibly due to a decrease in animal health following the physical stress of the injections (p = 0.79, WRS, Figure 4E). After 4-AP, Shaker RNAi-treated animals had weaker responses to coherent looms than did wild-type ones (compare Figure 3E and 4E; p = 0.030, WRS) and the 4-AP-dependent increase in firing was slightly lower than in wild-type animals for all black looming and translating visual stimuli, suggesting that the protein encoded by the *Shaker* gene is in part responsible for the 4-AP-sensitive conductance in field A (compare Figure 4F and G).

### The 4-AP-sensitive conductance in the LGMD is slowly-inactivating

When the LGMD dendritic field A was voltage clamped at a hyperpolarizing potential, then stepped to a depolarizing one, the result was an outward current that quickly rose, then slowly decayed back down to a non-zero steady state for sufficiently large depolarization steps (Figure 5A). After adding 4-AP, however, the same protocol resulted in a smaller outward current that slowly rose to a steady state (Figure 5B). The difference between those two currents (i.e., the 4-AP sensitive component) had a quick rise and slow decay, the characteristic shape of an inactivating voltage-gated ion channel (Figure 5C). To quantify the voltage dependence of activation as well as the activation and inactivation time constants, we performed the voltage clamp step protocol described above, keeping the pre-step hyperpolarizing potential constant at-85 mV while varying the depolarizing step (-70 to-20 mV; Figure 5D). For steady-state inactivation voltage dependence, the opposite protocol was used: the pre-step was varied (-100 mv to-60 mV) while the depolarizing step was held constant at-35 mV (Figure 5E-H).

**Figure 5.**
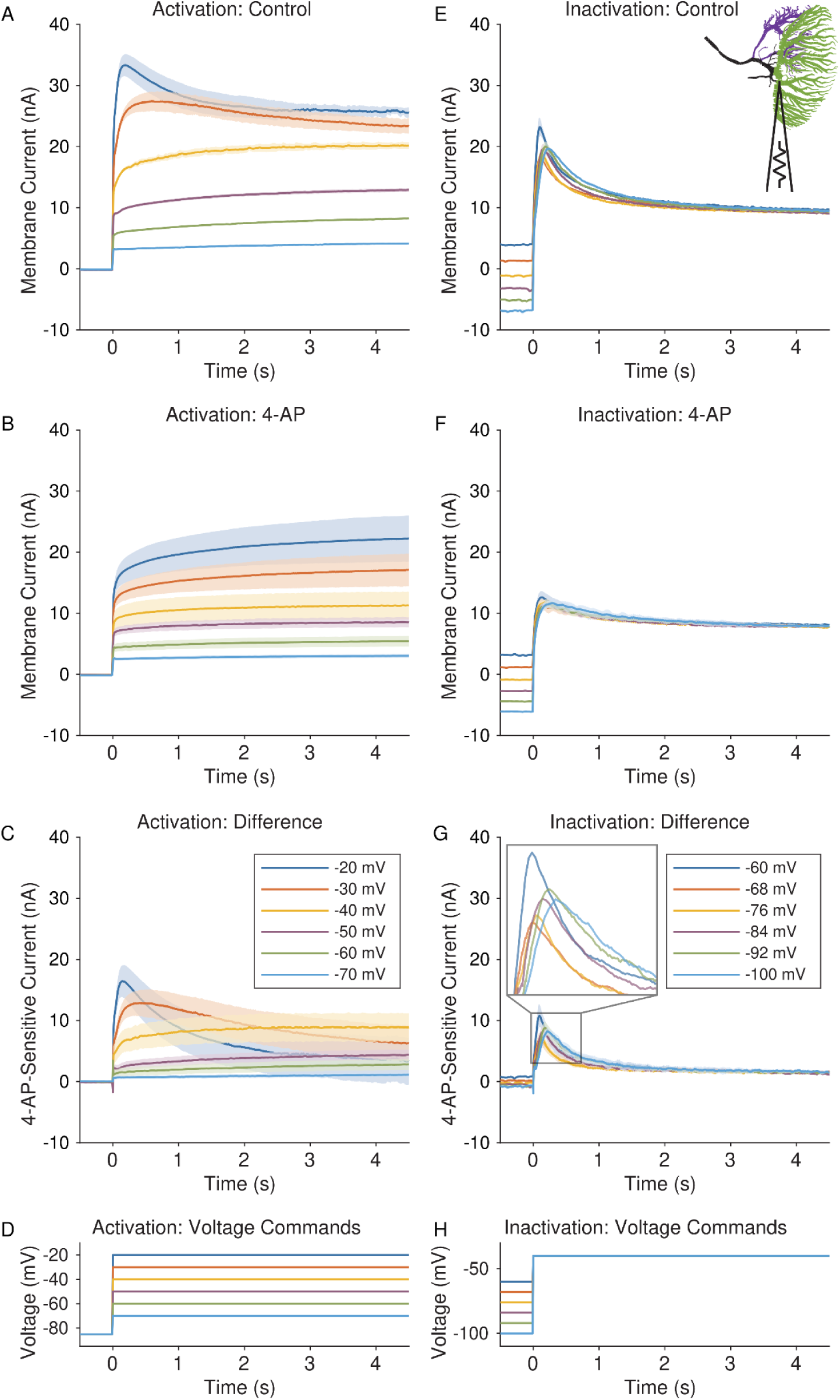
The 4-AP-sensitive conductance is a voltage-gated, slowly inactivating potassium conductance. A) Current response to activation characterization voltage clamp steps in the control condition, mean ± s.d. for one animal. All steps start at-85 mV before stepping to a variable depolarized potential. B) Same as A, after application of 4-AP. C) Differences between mean control (A) and 4-AP (B) current responses for each activation step. D) Activation characterization voltage command protocol for each step in 5A-C. E) Current response to inactivation characterization voltage clamp steps in the control condition, mean ± s.d. for one animal. Trials start at a variable pre-step potential before stepping to-30 mV. F) Same as B, after application of 4-AP. G) Same as D, for each inactivation step response. Inset is a 3x magnification of the boxed region, displaying the peak region of the mean responses. H) Inactivation characterization voltage command protocol for each step in 5E-G. Figure top right: LGMD with approximate recording location.

Increasing the depolarization step magnitude caused a larger peak difference current with faster rising and falling phases, resulting in an earlier peak relative to the start of the voltage step. In contrast, while lowering the hyperpolarizing pre-step increased the magnitude of the peak current for most pre-step potentials, peak current was sometimes higher and earlier for the most depolarized pre-step and lower and later for the most hyperpolarized one (Figure 5F).

### The 4-AP-sensitive conductance is active at behaviorally relevant potentials and timescales

By fitting subtractive 4-AP-sensitive conductance data, we obtained the half maximum voltage for activation and inactivation (-54 ± 11 mV and-79 ± 4 mV, mean ± s.d., n = 12 and 5 respectively) and the percentage permissiveness (8.5% and 40.8% respectively, Figure 6A) at the LGMD’s resting membrane potential (-65 mV). For the purpose of these calculations, the reversal potential of potassium was estimated at-75 mV, since the steady-state of the 4-AP-sensitive current was always positive at-70 mV and negative at-80 mV. Depolarization in field A during the response to a looming stimulus would activate a large fraction of the conductance, and higher input resistance in the thinner branches of field A means that they reach even higher voltages. Hence the conductance would play an even greater role in shaping responses in the cell’s periphery. This suggests that the 4-AP-sensitive conductance is active during peak depolarization (and therefore peak firing) of a loom response.

**Figure 6.**
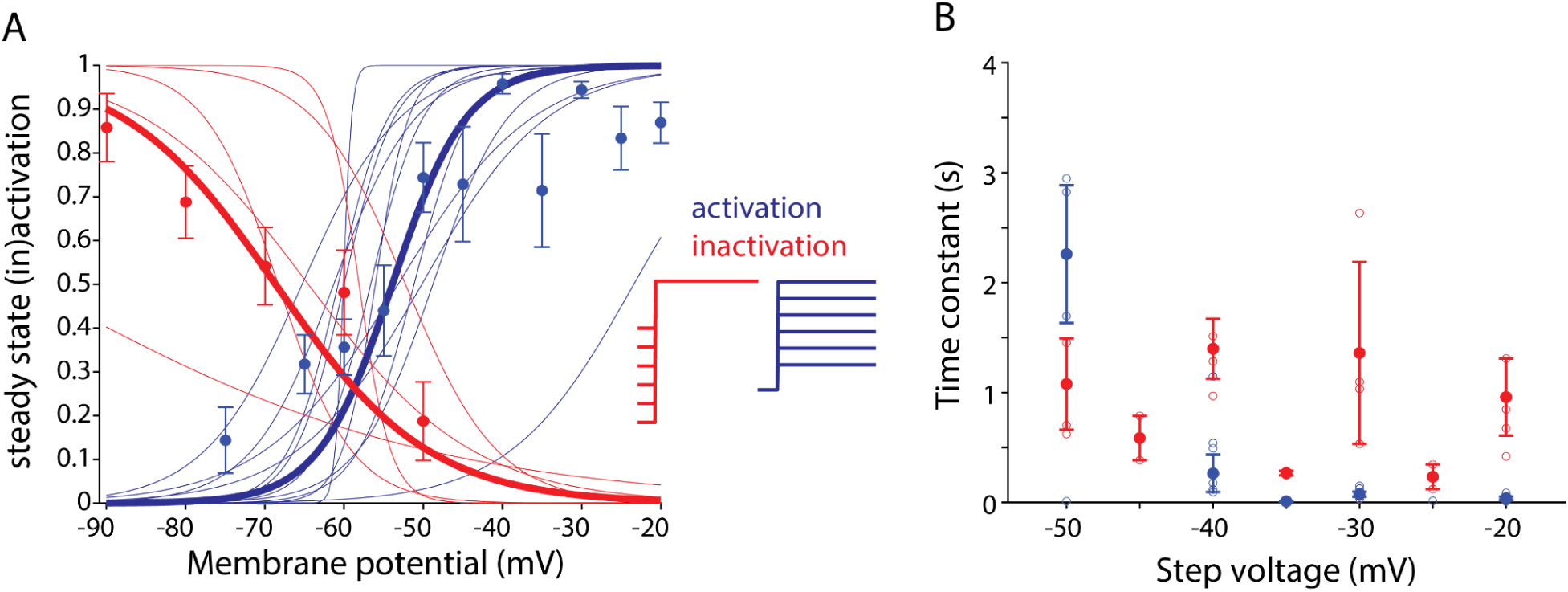
The 4-AP-sensitive conductance has voltage dependence and time constants in physiologically relevant ranges. A) Plot of percentage of 4-AP-sensitive channel activation at each of the activation voltages in 5D (blue) and pre-step inactivation voltages in 5H (red). Circles represent the mean across animals ± s.e.m. Thin curves are fit to individual animal means, while the thick curve is a fit to all of those individual means for all animals combined. B) Activation (blue) and inactivation (red) time constants at each voltage where rates were measurable. Solid circles represent the median across animals ± median deviation, open circles are individual animal values.

The activation time constant is on the order of tens of milliseconds at the voltages that the base of field A reaches during coherent looming stimuli (Figure 6B; Dewell and Gabbiani 2019). In the same voltage range, the inactivation time constant was on the order of hundreds of milliseconds to seconds, a similar timescale to that of looming stimuli (Figure 6B, Figure 3). Both time constants were faster at higher activation voltages.

### RNAi knock-down of Shaker increases LGMD intrinsic excitability and reduces the effect of 4-AP

Knocking down the *Shaker* channel gene (which, as shown in Figure 4, changed baseline visual responses and reduced their sensitivity to 4-AP) also resulted in changes to the LGMD’s electrical excitability and reduced the effect of 4-AP application. A depolarizing step of +6 nA to the base of LGMD field A results in high initial firing that decays over time (Figure 7A). At steady-state, towards the end of a 3-second step, firing drops to almost zero in wild-type animals, but 4-AP increases firing in that period (p = 0.063, WSR, Figure 7B). 4-AP also caused an increase in peak firing frequency (though not the timing of the peak) and in firing rate a fixed period after the peak (see Supplemental Figure 5). Shaker RNAi animals had higher baseline steady-state firing (p = 0.032 compared to wild-type control, WRS, Figure 7B), but firing after 4-AP was about the same as wild-type (p = 0.063 compared to Shaker RNAi control; p = 0.84 compared to wild-type 4-AP, WRS; Figure 7B).

**Figure 7.**
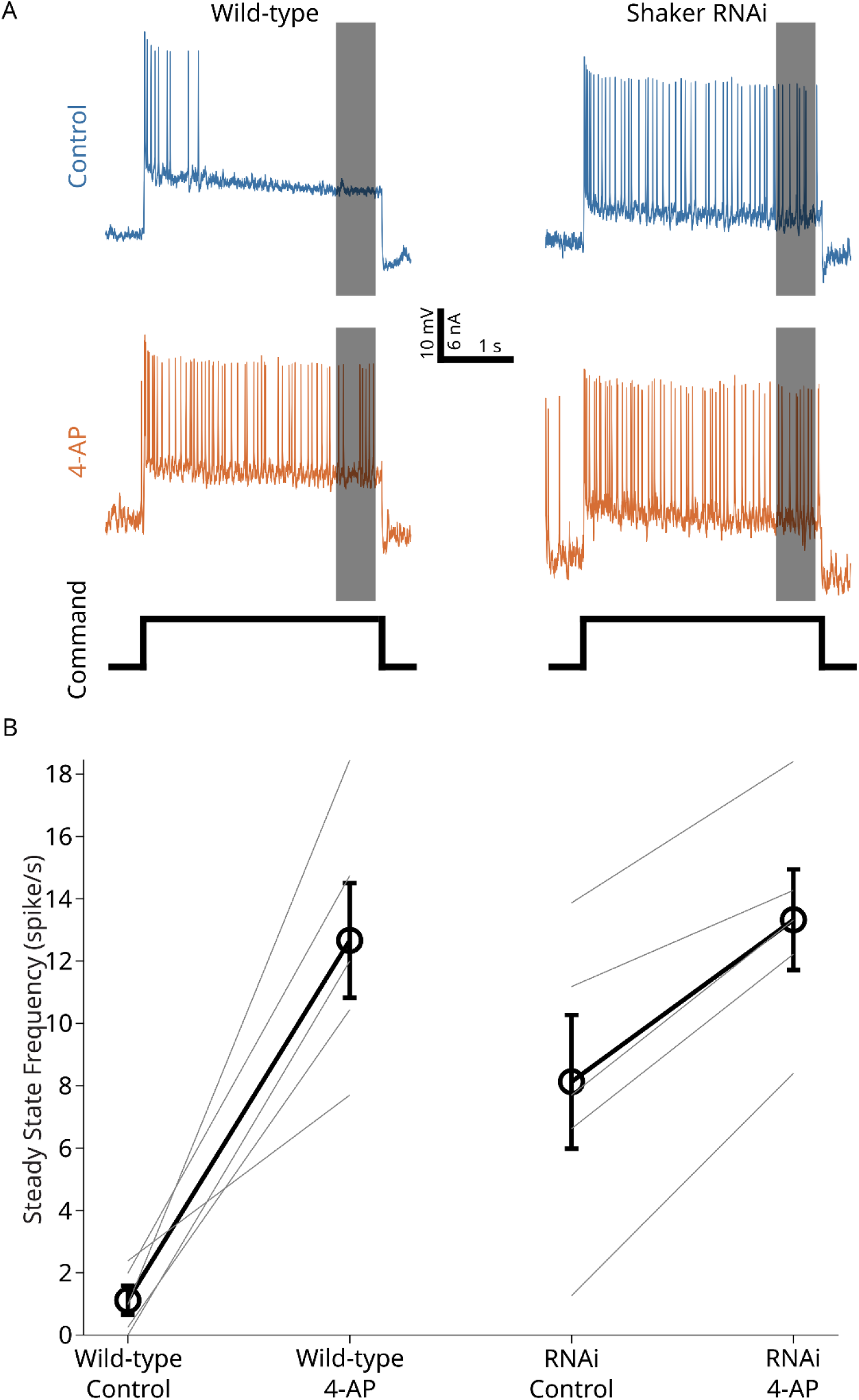
The LGMD in Shaker RNAi animals is more excitable and less 4-AP-sensitive. A) Voltage response to a +6 nA current step. Example voltage traces for a WT animal (left) and a Shaker RNAi animal (right) before (blue) and after 4-AP (orange). Current step protocol shown at the bottom in black. Grey boxes indicate the time period used to calculate steady-state firing below. B) Steady-state firing frequency during the last 0.5 s (the grey boxes in A) of a 3 s, +6 nA current step in WT and Shaker RNAi animals before and after 4-AP. Thick lines represent mean across animals ± s.e.m. (n = 5 each condition), with individual animals represented by thin lines.

### NEURON simulations that include experimentally-determined channel kinetics recapitulate real LGMD response properties

To test the effect of the newly characterized conductance and quantify its accuracy in recapitulating experimental results, we updated the existing multi-compartment functional model of the LGMD (Model DB #256024) with experimentally determined channel kinetics. For each of the four looming stimuli tested (black or white, 0% or 100% coherence), we created a simulation of the corresponding pattern and time course of synaptic inputs onto the LGMD’s dendritic tree, based on previous results that included imaging calcium fluorescence in the LGMD during single-facet stimulation (Zhu and Gabbiani 2016). We measured both the resulting firing pattern and subthreshold activity of the LGMD in the model (Figure 8). When the model includes a slowly-inactivating voltage-gated potassium conductance with experimentally-determined kinetics, the model reproduces results observed in experimental data, such as the preference for spatially coherent black stimuli. Without that conductance, results are less similar to control data and include phenomena that are more consistent with the LGMD after application of 4-AP, such as increased firing in the early stages of a looming stimulus response.

**Figure 8.**
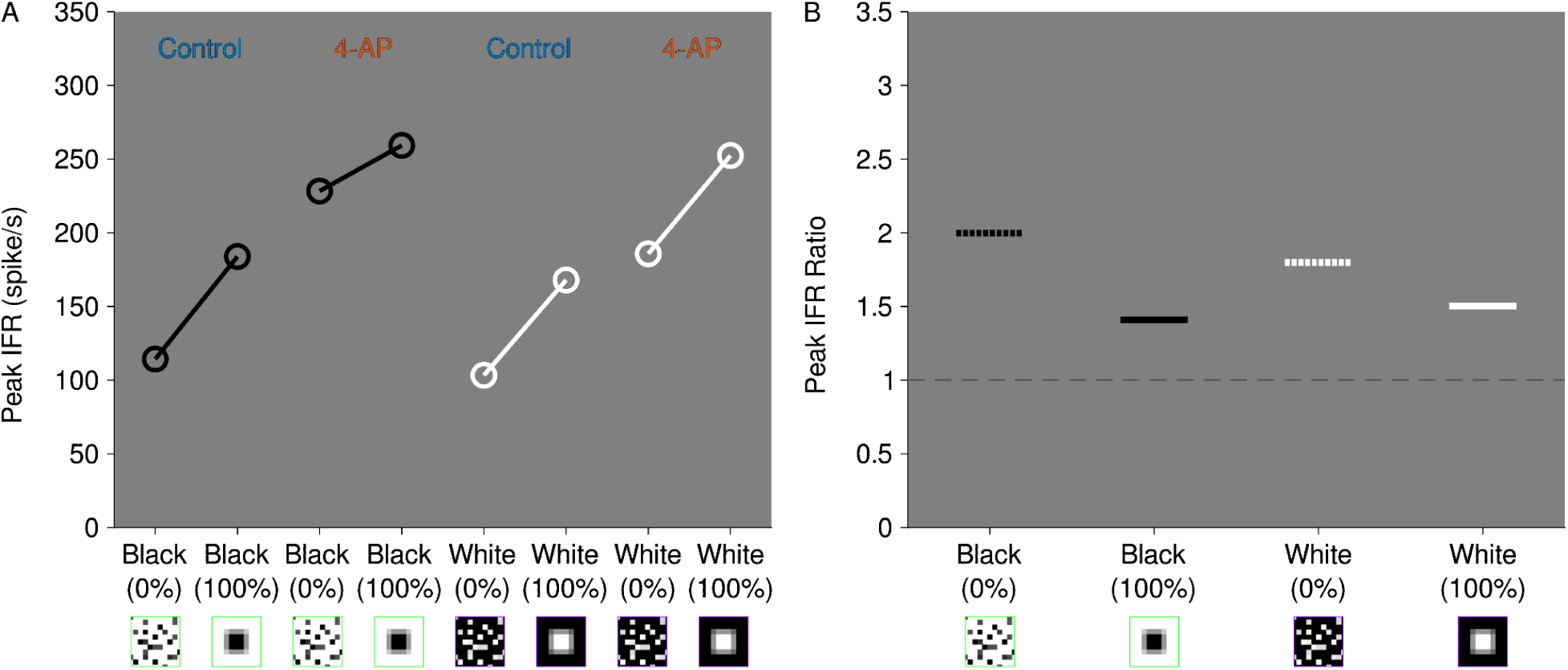
NEURON simulations of the LGMD reproduce trends in experimental responses to visual and electrical stimuli when they include an inactivating potassium channel in field A. A) Peak IFRs from simulated LGMD responses to the stimuli in Figure 3. 4-AP simulated by removing 95% of the 4-AP-sensitive conductance. B) Ratios in the peak IFRs in A. Dotted line represents a ratio of 1 (i.e., no change).

## Discussion

### *S. americana* Shaker produces the 4-AP-sensitive conductance

A high-quality, annotated genome of *Schistocerca americana* was, until recently, one missing piece in the puzzle of LGMD structure-function relationship. Here, the assembly iqSchAmer2.1 has been used in combination with single-cell RNA sequencing from LGMD cell bodies to identify K^+^ channel encoding genes and their relative levels of expression. In other species, these channels exhibit inactivation across a range of voltages and time scales (Wei et al. 1990). Some of these candidates normally generate channels with properties that are outside the range of observed voltage dependence and time constants of the 4-AP-sensitive conductance in the LGMD, although both properties vary with sequence changes (Wei et al. 1990; Imbrici et al. 2006; Waters et al. 2006), RNA editing (Bhalla et al. 2004), post-translational modifications (Jindal et al., 2008), and the presence of protein co-factors such as β subunits (Morales et al. 1995; Schulte et al. 2006). The most highly expressed voltage-gated potassium channel in the LGMD was a Shaker channel that has the conserved 4-AP binding site (Pinto-Anwandter 2024). The detection of DLG in the LGMD is also of interest, as it could regulate the observed subcellular localization of the 4-AP-sensitive conductance.

Phylogenetic analysis of the *Shaker* family of voltage-gated potassium channels (*Shaker*, *Shab*, *Shaw*, and *Shal*, corresponding to mammalian channel genes KCNA, KCNB, KCNC, and KCND, respectively) revealed clustering of genes from *Schistocerca*, other arthropods, and animals as genetically distant as the freshwater hydra. Despite at least half a billion years of evolution between *Schistocerca americana* and *Hydra vulgaris*, the grasshopper *Shaker* is more similar to that of hydra than it is to *Shaw* (Figure 2B; Fedonkin et al. 2007).

Sequence features like the 4-AP binding site were conserved (Figure 2B). The considered species process sensory information across a wide range of environments and time scales using essentially the same set of ion channels (Ruck 1961; Healy et al. 2013; Petie et al. 2016). This suggests that the ability of neurons to respond appropriately to sensory stimuli depends on additional factors, such as neuron morphology, synaptic connectivity, expression patterns, and channel modifications.

### LGMD responses to behaviorally-irrelevant stimuli are preferentially suppressed by a 4-AP-sensitive potassium conductance in field A

The effect of 4-AP on LGMD visual responses presented here builds on work which showed that 4-AP causes an increase in firing inversely related to looming stimulus coherence for black stimuli (Dewell and Gabbiani 2018). Thus, under normal conditions the 4-AP-sensitive conductance selectively suppresses responses to incoherent black stimuli (Figure 9).

**Figure 9.**
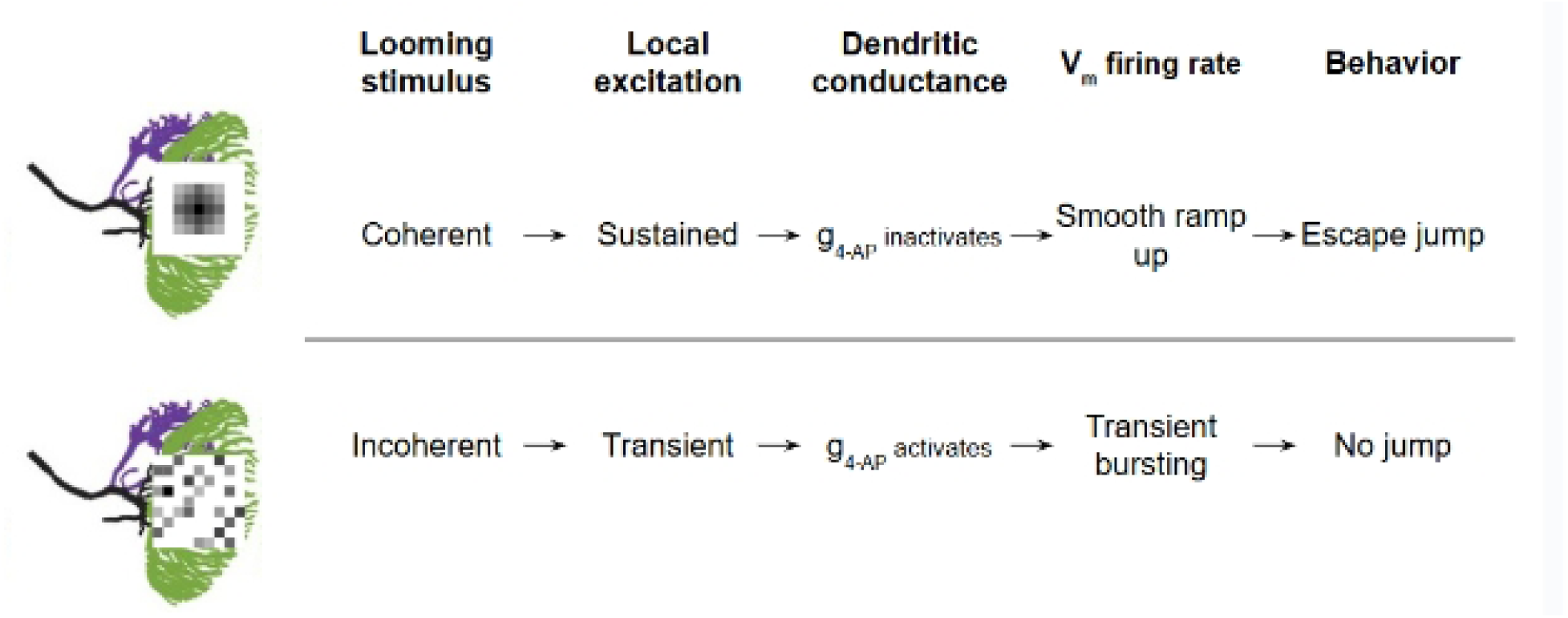
Physiological and behavioral effects of coherence selectivity. For a solid loom, the sequential activation of many neighboring facets causes the corresponding part of the LGMD to have a large depolarization that lasts long enough for this conductance to inactivate. With lower potassium conductance, Vm is able to ramp up smoothly to a high peak, resulting in the correct firing pattern and a jump. During a scrambled loom, however, individual inputs are spread out and cannot summate, meaning that EPSPs come in and quickly decay, lasting long enough to activate the conductance (further reducing their strength) but not long enough to inactivate it. These weak, transient depolarizations produce a lower, choppier firing rate that is insufficient to produce a jump.

Conversely, an HCN conductance potentiates responses to coherent stimuli, and together the two establish the LGMD’s coherence preference for black looming stimuli (Dewell and Gabbiani 2018). In the LGMD, only field A exhibits retinotopy (Zhu and Gabbiani 2016; Dewell et al. 2022). Since spatial coherence detection requires retinotopy, there would be no use for the expression of a coherence-detecting conductance in a non-retinotopic area. Indeed, application of 4-AP to field C yielded no changes in visual responses. This is consistent with the predicted absence of the 4-AP-sensitive conductance in field C and suggests that its dendritic expression is related to coherence discrimination.

### The 4-AP-sensitive conductance shapes visual responses at behaviorally relevant potentials and timescales

Based on the calculated g_4-AP_ kinetics, the steady-state voltage dependencies of both activation and inactivation gates predict a low-conductance state while the cell is at rest, so small voltage changes cause relatively large changes in percent conductance. During visual stimulation, excitatory inputs cause an increase in 4-AP-sensitive conductance that decreases input resistance and dendritic electrotonic length, preventing the summation of excitatory inputs.

During a coherent looming stimulus, however, retinotopy causes localized, clustered excitation, allowing depolarization to reach a level where the 4-AP-sensitive conductance shuts off due to inactivation. This suggests that this conductance preferentially suppresses short, discrete excitatory inputs but not large, sustained ones during visual stimulation.

### Knocking down Shaker partially recapitulates 4-AP effects

To the best of our knowledge, this is the first instance of RNAi in *Schistocerca* characterized by *in vivo* intracellular recordings of a neuronal phenotype. If the channel encoded by the *Shaker* gene underlies the 4-AP-sensitive conductance, one expects that knocking it down would result in a phenotype that resembles the effect of 4-AP, and that further 4-AP application would have little effect. This prediction was partially borne out since qPCR and physiological results showed that RNAi reduced but did not entirely eliminate the targeted conductance. This could be because not all Shaker mRNA was eliminated, or because the 4-AP-sensitive conductance includes non-Shaker channels. Another possibility is that knocking-down of Shaker for over a week caused a compensatory change, such as another channel being upregulated, that did not occur during the hour-long 4-AP application. Additional systematic efforts to titrate the dsRNA dose, incubation time, or number of boosters may provide information about the timing and regulation of Shaker channel expression in the LGMD. Indeed, the remaining 4-AP-sensitive conductance may result from the delay between transcription and translation, low channel turnover, or a reserve pool of channel proteins. Even if an antibody allowed for a quantitative analysis of total Shaker protein levels in bulk tissue, it would not provide a direct measure of reduction in functional channels since post-translational modification or insertion into the LGMD membrane could be limiting factors.

The strengthening effect of both 4-AP and Shaker RNAi on sustained firing in response to sustained depolarization demonstrates that the conductance plays a role in spike frequency adaptation (SFA). Calcium-dependent potassium “small-conductance” (SK) channels have been implicated in regulating SFA in the LGMD (Peron and Gabbiani 2009). The contribution of the slowly-inactivating 4-AP-sensitive conductance suggests that SFA in the LGMD involves both calcium-dependent and-independent mechanisms.

### Simulations of the LGMD demonstrate how ion channel expression shapes neural responses

The NEURON model of the LGMD allows rapid simulation of visual or intracellular experiments and recording of parameters that are inaccessible *in-vivo*. In the model, we can measure voltage changes in every part of the cell simultaneously such as the rapid voltage attenuation of early inputs during an incoherent looming stimulus. We can also directly measure how individual conductances change, such as the greater drop in the 4-AP-sensitive conductance during a coherent loom. One limitation is that neurons immediately upstream of the LGMD were not modeled, so the strength of synaptic inputs was set by comparison to experimental results (Jones and Gabbiani 2012). In addition, each conductance in the LGMD, including the 4-AP-sensitive one, is represented by a single, uniform channel type. In the real cell, channels may be heteromeric or differentially modified by RNA editing, post-translational modification, and auxiliary subunits. This would mean that the experimentally observed conductance may be the result of a mix of channels with different voltage dependences or time constants.

### Role of a slowly-inactivating voltage-gated potassium conductance in spatial coherence discrimination

We have shown that, in the LGMD, a *Shaker* gene contributes to accurate predator detection by selectively weakening responses to looming stimuli according to spatial coherence. Our neuronal modeling approach demonstrates that it is possible to characterize the contribution of an individual conductance to complex neuronal computations. From this, it can be inferred that similar computations are likely being performed by neurons in the brains of other organisms, including our own.

## Acknowledgements

Thanks to Dr. Saina Namazifard for advice on fitting curves to current traces. Thanks to Dr. Edward Cooper for advice on channel phylogeny.

## Figure Code

Matlab code and data to generate figures can be found at https://data.mendeley.com/preview/5gkgkmphhc?a=8700d09d-397b-48e9-8802-6f3f91b44b63

## Supplement

**Supplemental Figure 1:**
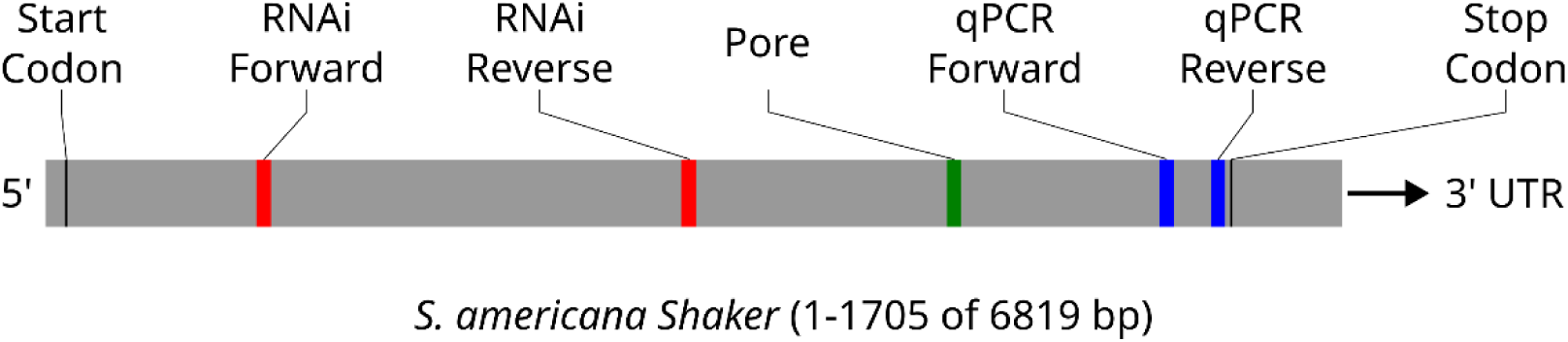
Map of *Shaker* mRNA with labeled features, to scale, including the start and stop codons, forward and reverse primers for both RNAi and qPCR, and the six codons of the pore region (TTVGYG). Most of the 3’ untranslated region (UTR) is omitted for concision.

**Supplemental Figure 2:**
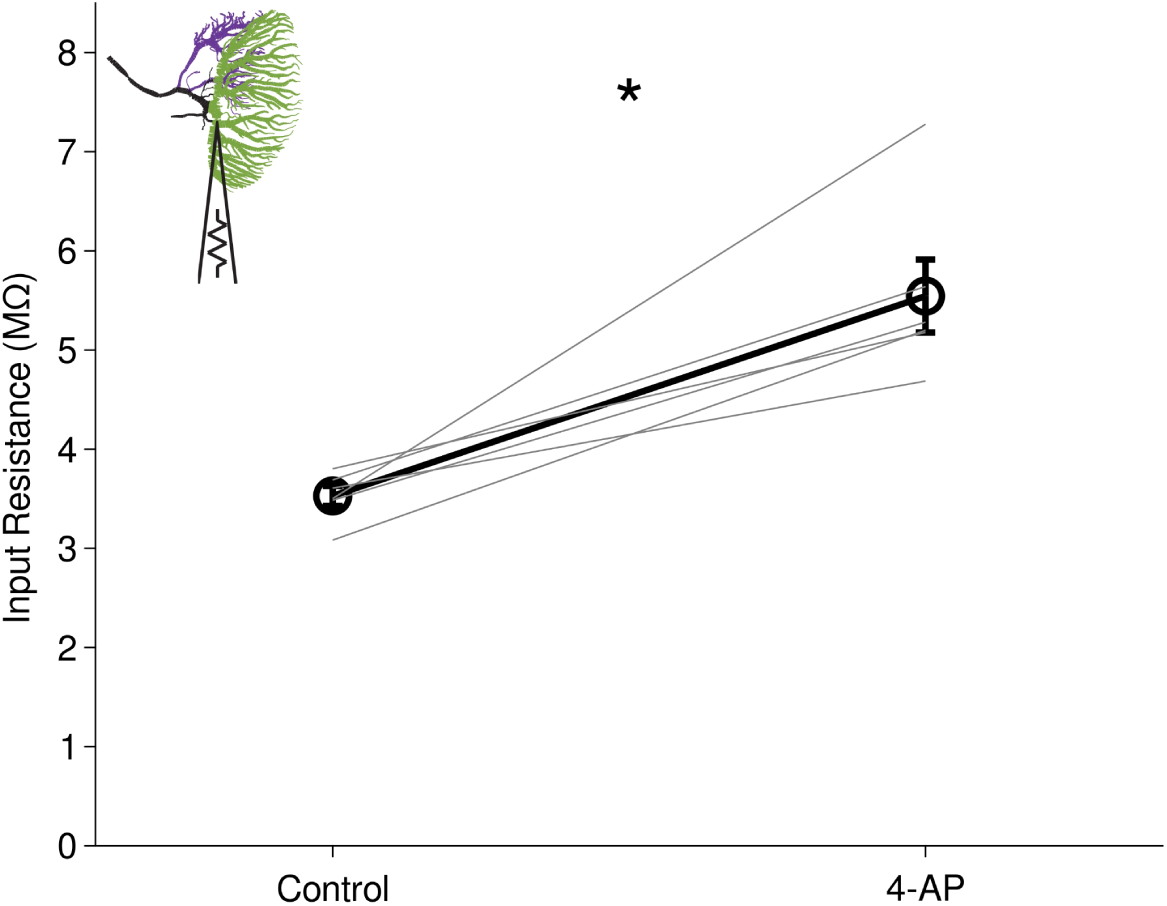
LGMD Input resistance at the base of field A, calculated from a-2 nA step in current clamp, before and after 4-AP, n = 6. Thick lines are mean ± s.e.m. across animals, thin lines are individual animals. Input resistance increased from 3.5 ± 0.1 MΩ to 5.5 ± 0.4 MΩ (p = 0.031, WSR).

**Supplemental Figure 3:**
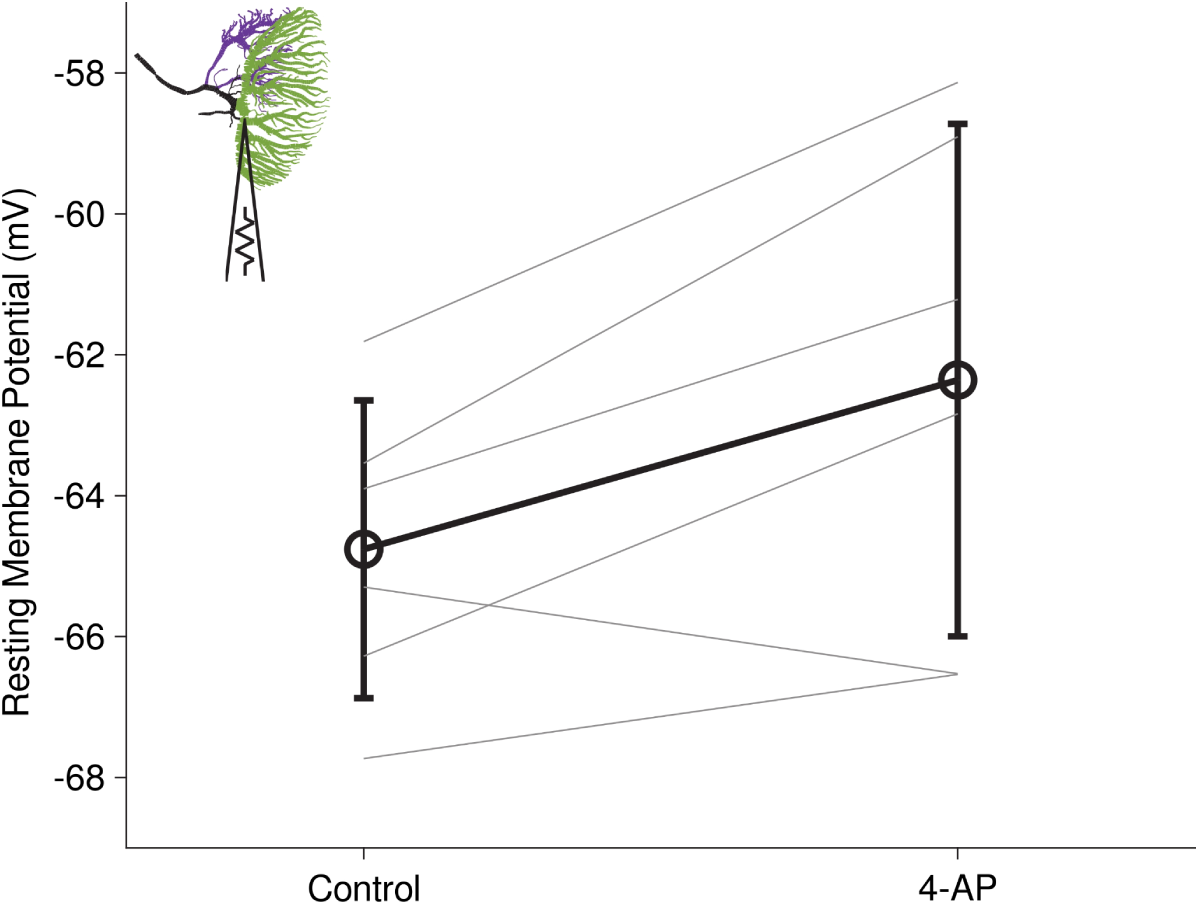
LGMD resting membrane potential at the base of field A, before and after 4-AP, n = 6. Black lines are mean ± s.e.m. across animals, gray lines are individual animals. Resting membrane potential increased from-64.8 ± 0.9 mV to-62.4 ± 1.5 mV (p = 0.094, WSR).

**Supplemental Figure 4:**
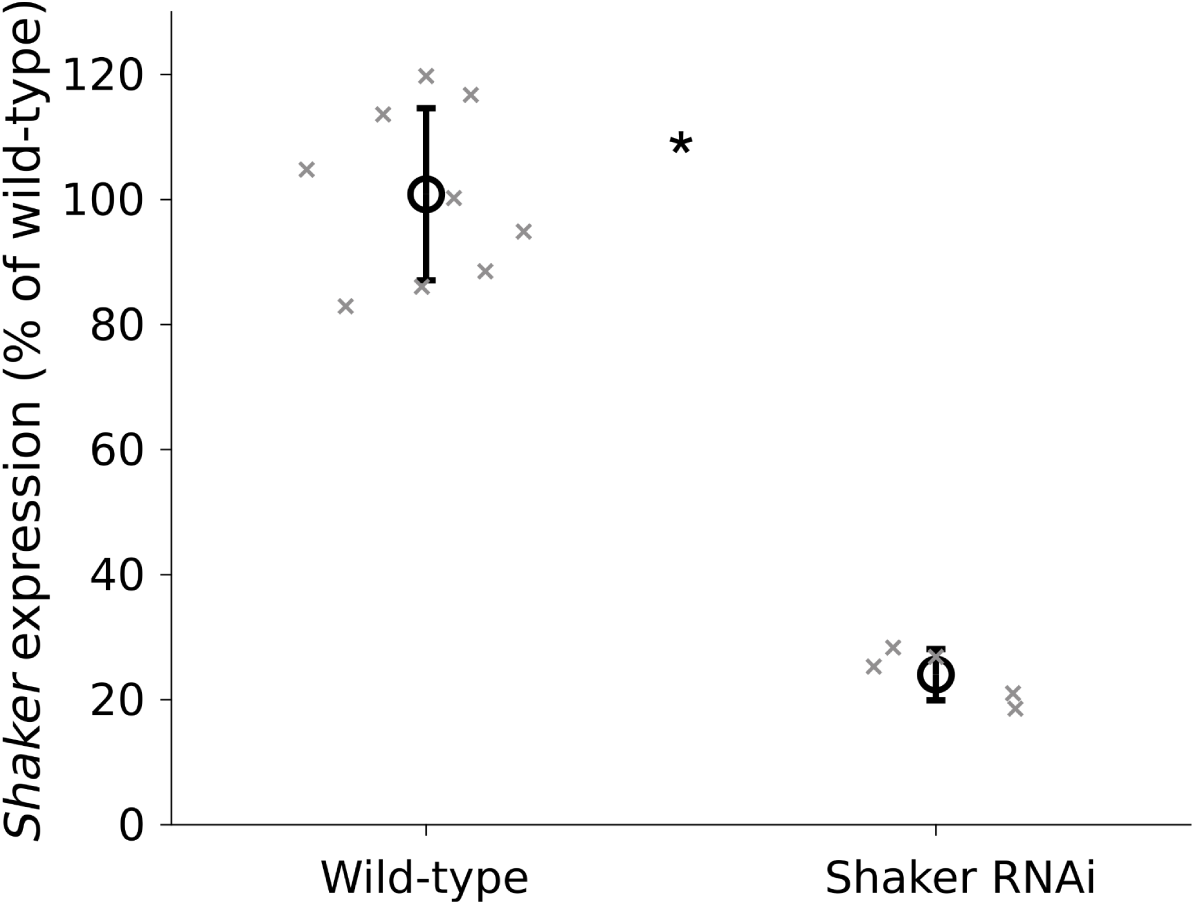
*Shaker* RNAi efficacy. Percent expression levels of *Shaker* mRNA in wild-type and *Shaker*-RNAi animals, relative to wild-type average. Black lines are mean ± s.d. across animals, gray marks are individual animals (jittered to improve visibility; p = 0.0001, WRS).

**Supplemental Figure 5:**
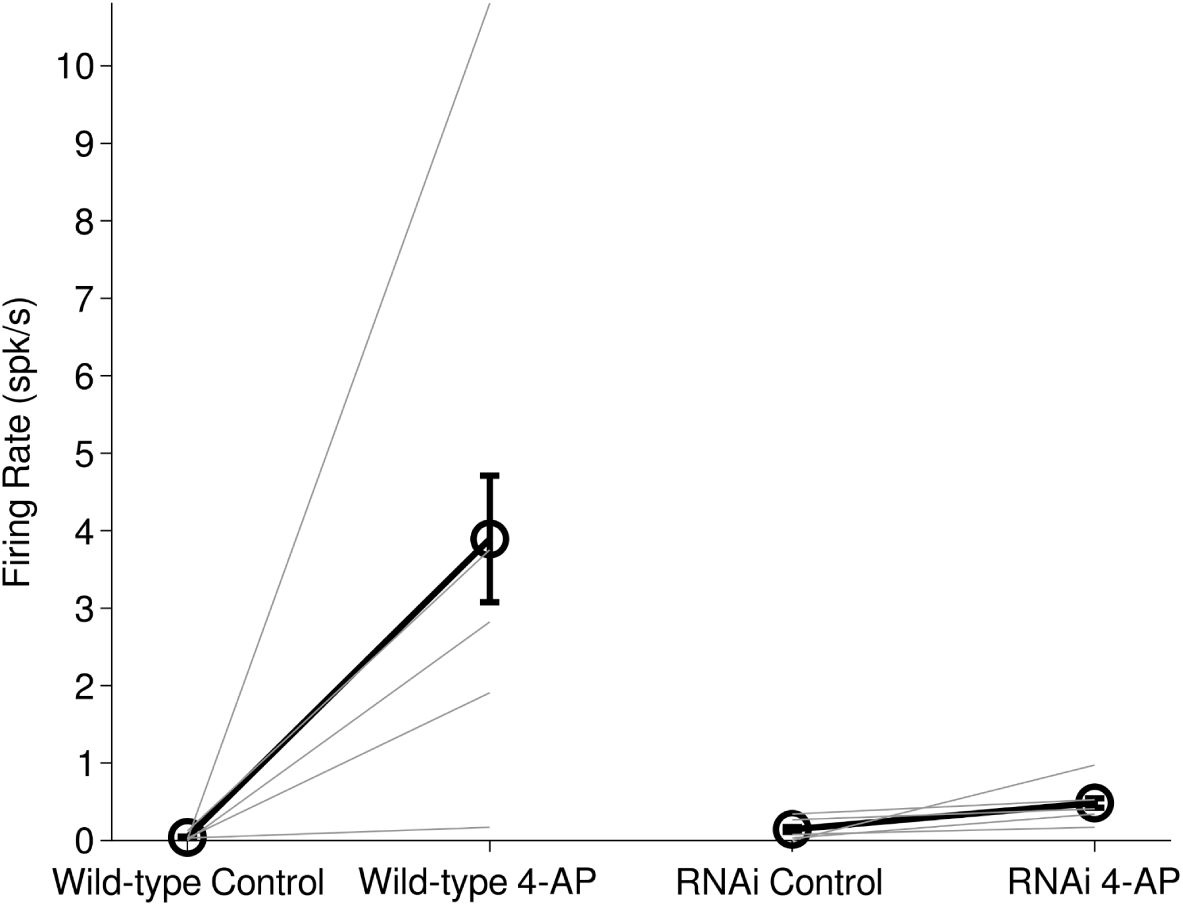
Spontaneous firing frequency, WT and Shaker RNAi animals, n = 5 each, ± 4-AP. Thick lines are mean ± s.e.m. across animals, thin lines are individual animals. 4-AP increased spontaneous firing relative to control in both wild-type and Shaker-RNAi animals (both p = 0.063, WSR).

**Supplemental Figure 6:**
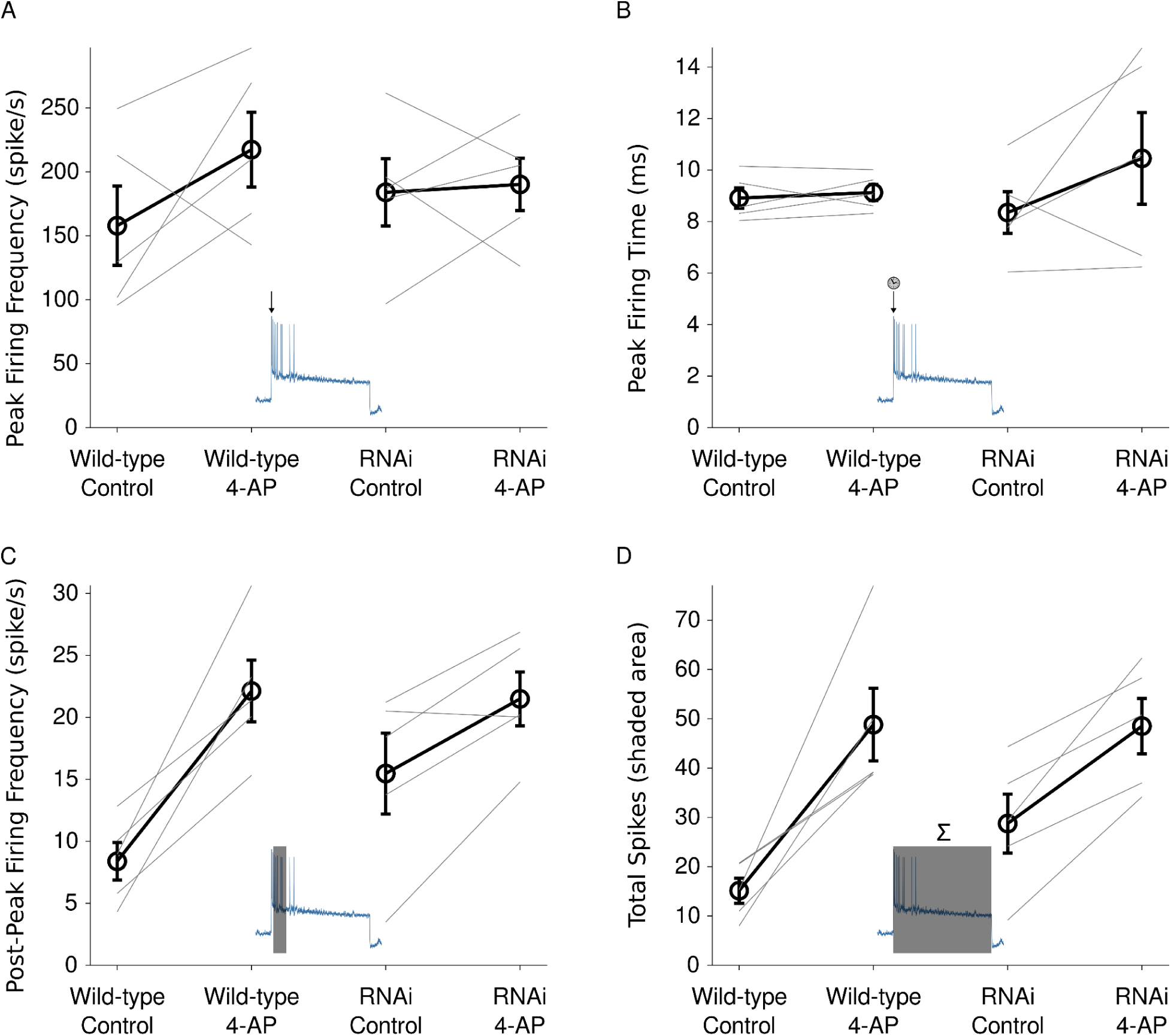
Firing properties in response to a 3-second +6 nA current step in wild-type and Shaker RNAi animals before and after 4-AP. Black lines represent mean across animals ± s.e.m, with individual animals represented by gray lines. A) Peak firing frequency (spike/s). B) Time of peak firing (ms). C) Firing frequency during a fixed period, between 100 ms and 500 ms, after the time of peak firing, (spike/s). D) Total number of spikes during step.

---

**Supplemental Table 1:** LGMD expression levels of potassium channel transcripts and related genes from single-cell mRNA sequencing, n = 5.

| Gene ID | *S. americana*<br>gene | Mammalian<br>gene family | Mammalian<br>Protein Homolog | CPM mean | CPM s.e.m. |
| --- | --- | --- | --- | --- | --- |
| LOC124612295 | *Shaker* | *KCNA* | Kv1 | 5,617.65 | 1,238.81 |
| LOC124606852 | *Shab* | *KCNB* | Kv2 | 208.94 | 25.10 |
| LOC124595212 | *Shaw1* | *KCNC* | Kv3 | 505.65 | 80.30 |
| LOC124595754 | *Shaw2* | *KCNC* | Kv3 | 139.74 | 28.08 |
| LOC124556270 | *Shal* | *KCND* | Kv4 | 562.17 | 67.99 |
| LOC124550780 | *Eag* | *KCNH* | Kv10 | 449.70 | 74.80 |
| LOC124556243 | *Erg* | *KCNH* | Kv11 | 1,336.09 | 132.02 |
| LOC124624305 | *Elk* | *KCNH* | Kv12 | 157.85 | 22.65 |
| LOC124553489 | *Shaker-β* | *KCNAB* | Kvβ | 2,131.88 | 321.32 |
| LOC124596031 | *DLG* | *DLG* | PSD-93 | 869.19 | 130.56 |

**Supplemental Table 2:**
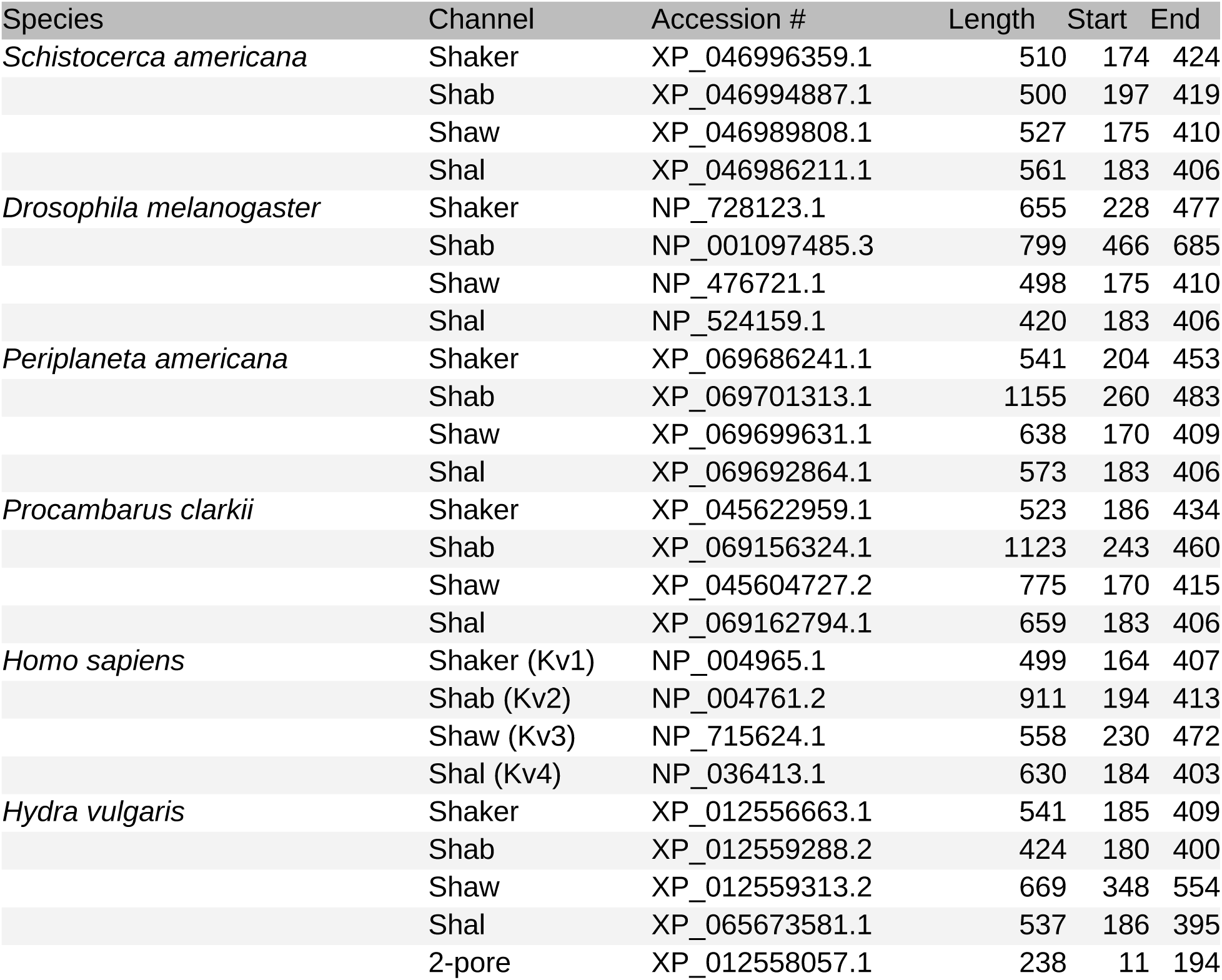
Homologous channels across species, including NCBI accession number, protein length, and the numbers of the first residue of the first transmembrane helix (Start) and the last residue of the last (End), which are the boundaries of the segments used to generate the phylogenetic tree in Figure 2A.

## References

Altschul, S. F., T. L. Madden, A. A. Schäffer, et al. 1997. “Gapped BLAST and PSI-BLAST: A New Generation of Protein Database Search Programs.” Nucleic Acids Research 25 (17): 3389–402. 10.1093/nar/25.17.3389.

Bhalla, Tarun, Joshua J. C. Rosenthal, Miguel Holmgren, and Robert Reenan. 2004. “Control of Human Potassium Channel Inactivation by Editing of a Small mRNA Hairpin.” Nature Structural & Molecular Biology 11 (10): 950–56. 10.1038/nsmb825.

Burrows, M., and C. H. F. Rowell. 1973. “Connections between Descending Visual Interneurons and Metathoracic Motoneurons in the Locust.” Journal of Comparative Physiology 85 (3): 221–34. 10.1007/BF00694231.

Curran, Mark E., Gregory M. Landes, and Mark T. Keating. 1992. “Molecular Cloning, Characterization, and Genomic Localization of a Human Potassium Channel Gene.” Genomics 12 (4): 729–37. 10.1016/0888-7543(92)90302-9.

Desper, Richard, and Olivier Gascuel. 2004. “Theoretical Foundation of the Balanced Minimum Evolution Method of Phylogenetic Inference and Its Relationship to Weighted Least-Squares Tree Fitting.” Molecular Biology and Evolution 21 (3): 587–98. 10.1093/molbev/msh049.

Dewell, Richard B., and Fabrizio Gabbiani. 2019. “Active Membrane Conductances and Morphology of a Collision Detection Neuron Broaden Its Impedance Profile and Improve Discrimination of Input Synchrony.” Journal of Neurophysiology 122 (2): 691–706. 10.1152/jn.00048.2019.

Dewell, Richard Burkett, and Fabrizio Gabbiani. 2018. “Biophysics of Object Segmentation in a Collision-Detecting Neuron.” eLife 7 (April): e34238. 10.7554/eLife.34238.

Dewell, Richard Burkett, Ying Zhu, Margaret Eisenbrandt, Richard Morse, and Fabrizio Gabbiani. 2022. “Contrast Polarity-Specific Mapping Improves Efficiency of Neuronal Computation for Collision Detection.” eLife 11 (October): e79772. 10.7554/eLife.79772.

Dobin, Alexander, Carrie A. Davis, Felix Schlesinger, et al. 2013. “STAR: Ultrafast Universal RNA-Seq Aligner.” Bioinformatics 29 (1): 15–21. 10.1093/bioinformatics/bts635.

Fedonkin, M. A., A. Simonetta, and A. Y. Ivantsov. 2007. “New Data on Kimberella, the Vendian Mollusc-like Organism (White Sea Region, Russia): Palaeoecological and Evolutionary Implications.” In The Rise and Fall of the Ediacaran Biota, edited by Patricia Vickers-Rich and Patricia Komarower, vol. 286. Geological Society of London. 10.1144/SP286.12.

Fotowat, Haleh, and Fabrizio Gabbiani. 2007. “Relationship between the Phases of Sensory and Motor Activity during a Looming-Evoked Multistage Escape Behavior.” The Journal of Neuroscience 27 (37): 10047–59. 10.1523/jneurosci.1515-07.2007.

Fotowat, Haleh, and Fabrizio Gabbiani. 2011. “Collision Detection as a Model for Sensory-Motor Integration.” Annual Review of Neuroscience 34: 1–19. 10.1146/annurev-neuro-061010-113632.

Fotowat, Haleh, Reid R. Harrison, and Fabrizio Gabbiani. 2011. “Multiplexing of Motor Information in the Discharge of a Collision Detecting Neuron during Escape Behaviors.” Neuron 69 (1): 147–58. 10.1016/j.neuron.2010.12.007.

Gabbiani, Fabrizio, Holger G. Krapp, and Gilles Laurent. 1999. “Computation of Object Approach by a Wide-Field, Motion-Sensitive Neuron.” ARTICLE. Journal of Neuroscience 19 (3): 1122–41. 10.1523/JNEUROSCI.19-03-01122.1999.

Grishin, Nick V. 1995. “Estimation of the Number of Amino Acid Substitutions per Site When the Substitution Rate Varies among Sites.” Journal of Molecular Evolution 41 (5): 675–79. 10.1007/BF00175826.

Gür, Burak, Katja Sporar, Anne Lopez-Behling, and Marion Silies. 2020. “Distinct Expression of Potassium Channels Regulates Visual Response Properties of Lamina Neurons in Drosophila Melanogaster.” Journal of Comparative Physiology A 206 (2): 273–87. 10.1007/s00359-019-01385-7.

Hallgren, Jeppe, Konstantinos D. Tsirigos, Mads Damgaard Pedersen, et al. 2022. “DeepTMHMM Predicts Alpha and Beta Transmembrane Proteins Using Deep Neural Networks.” Preprint, bioRxiv, April 10. 10.1101/2022.04.08.487609.

Healy, Kevin, Luke McNally, Graeme D. Ruxton, Natalie Cooper, and Andrew L. Jackson. 2013. “Metabolic Rate and Body Size Are Linked with Perception of Temporal Information.” Animal Behaviour 86 (4): 685–96. 10.1016/j.anbehav.2013.06.018.

Hodgkin, A. L., and A. F. Huxley. 1952. “A Quantitative Description of Membrane Current and Its Application to Conduction and Excitation in Nerve.” The Journal of Physiology 117 (4): 500–544. 10.1113/jphysiol.1952.sp004764.

Hoffman, Dax A., Jeffrey C. Magee, Costa M. Colbert, and Daniel Johnston. 1997. “K+ Channel Regulation of Signal Propagation in Dendrites of Hippocampal Pyramidal Neurons.” Nature 387 (6636): 869–75. 10.1038/43119.

Imbrici, Paola, Maria Cristina D’Adamo, Dimitri M. Kullmann, and Mauro Pessia. 2006. “Episodic Ataxia Type 1 Mutations in the KCNA1 Gene Impair the Fast Inactivation Properties of the Human Potassium Channels Kv1.4-1.1/Kvβ1.1 and Kv1.4-1.1/Kvβ1.2.” European Journal of Neuroscience 24 (11): 3073–83. 10.1111/j.1460-9568.2006.05186.x.

Jacobs, Dave K., Nagayasu Nakanishi, David Yuan, Anthony Camara, Scott A. Nichols, and Volker Hartenstein. 2007. “Evolution of Sensory Structures in Basal Metazoa.” Integrative and Comparative Biology 47 (5): 712–23. 10.1093/icb/icm094.

Jan, Y. N., Lily Y. Jan, and M. J. Dennis. 1977. “Two Mutations of Synaptic Transmission in Drosophila.” Proceedings of the Royal Society of London. Series B. Biological Sciences 198 (1130): 87–108. 10.1098/rspb.1977.0087.

Jegla, Timothy, Christian Zmasek, Serge Batalov, and Surendra Nayak. 2009. “Evolution of the Human Ion Channel Set.” Combinatorial Chemistry & High Throughput Screening 12 (1): 2–23. 10.2174/138620709787047957.

Jones, Peter W., and Fabrizio Gabbiani. 2010. “Synchronized Neural Input Shapes Stimulus Selectivity in a Collision-Detecting Neuron.” Current Biology 20 (22): 2052–57. 10.1016/j.cub.2010.10.025.

Jones, Peter W., and Fabrizio Gabbiani. 2012. “Logarithmic Compression of Sensory Signals within the Dendritic Tree of a Collision-Sensitive Neuron.” Articles. Journal of Neuroscience 32 (14): 4923–34. 10.1523/JNEUROSCI.5777-11.2012.

Krapp, Holger G., and Fabrizio Gabbiani. 2005. “Spatial Distribution of Inputs and Local Receptive Field Properties of a Wide-Field, Looming Sensitive Neuron.” Journal of Neurophysiology 93 (4): 2240–53. 10.1152/jn.00965.2004.

Liao, Yang, Gordon K. Smyth, and Wei Shi. 2014. “featureCounts: An Efficient General Purpose Program for Assigning Sequence Reads to Genomic Features.” Bioinformatics 30 (7): 923–30. 10.1093/bioinformatics/btt656.

Mitra, Soumi, Saina Namazifard, David Mario Bellini, et al. 2025. “To Jump or Not to Jump: Comparing Effects of Phenotypic Plasticity on the Visual Responses and Escape Behavior of Locusts and Grasshoppers.” Preprint, bioRxiv, August 31. 10.1101/2025.08.26.672473.

Morales, Michael J., Robert C. Castellino, Anne L. Crews, Randall L. Rasmusson, and Harold C. Strauss. 1995. “A Novel β Subunit Increases Rate of Inactivation of Specific Voltage-Gated Potassium Channel α Subunits (∗).” Journal of Biological Chemistry 270 (11): 6272–77. 10.1074/jbc.270.11.6272.

Ogawa, Yasuhiro, Ido Horresh, James S. Trimmer, David S. Bredt, Elior Peles, and Matthew N. Rasband. 2008. “Postsynaptic Density-93 Clusters Kv1 Channels at Axon Initial Segments Independently of Caspr2.” The Journal of Neuroscience 28 (22): 5731–39. 10.1523/JNEUROSCI.4431-07.2008.

O’Shea, Michael, and J. L. D. Williams. 1974. “The Anatomy and Output Connection of a Locust Visual Interneurone; the Lobular Giant Movement Detector (LGMD) Neurone.” Journal of Comparative Physiology 91 (3): 257–66. 10.1007/BF00698057.

Peron, Simon, and Fabrizio Gabbiani. 2009. “Spike Frequency Adaptation Mediates Looming Stimulus Selectivity in a Collision-Detecting Neuron.” Nature Neuroscience 12 (3): 318–26. 10.1038/nn.2259.

Peron, Simon P., Peter W. Jones, and Fabrizio Gabbiani. 2009. “Precise Subcellular Input Retinotopy and Its Computational Consequences in an Identified Visual Interneuron.” Neuron 63 (6): 830–42. 10.1016/j.neuron.2009.09.010.

Petie, Ronald, Michael R. Hall, Mia Hyldahl, and Anders Garm. 2016. “Visual Orientation by the Crown-of-Thorns Starfish (Acanthaster Planci).” Coral Reefs 35 (4): 1139–50. 10.1007/s00338-016-1478-0.

Pinto-Anwandter, Bernardo I. 2024. “Structural Basis for Voltage Gating and Dalfampridine Binding in the Shaker Potassium Channel.” Preprint, bioRxiv, October 25. 10.1101/2024.10.22.619486.

Rettig, Jens, Stefan H. Heinemann, Frank Wunder, et al. 1994. “Inactivation Properties of Voltage-Gated K+ Channels Altered by Presence of β-Subunit.” Nature 369 (6478): 289–94. 10.1038/369289a0.

Robinson, Mark D., Davis J. McCarthy, and Gordon K. Smyth. 2010. “edgeR: A Bioconductor Package for Differential Expression Analysis of Digital Gene Expression Data.” Bioinformatics 26 (1): 139–40. 10.1093/bioinformatics/btp616.

Ruck, Philip. 1961. “Electrophysiology of the Insect Dorsal Ocellus.” The Journal of General Physiology 44 (3): 641–57. 10.1085/jgp.44.3.641.

Salkoff, Lawrence, Keith Baker, Alice Butler, Manuel Covarrubias, Michael D. Pak, and Aguan Wei. 1992. “An Essential ‘Set’ of K+ Channels Conserved in Flies, Mice and Humans.” Trends in Neurosciences 15 (5): 161–66. 10.1016/0166-2236(92)90165-5.

Schulte, Uwe, Jörg-Oliver Thumfart, Nikolaj Klöcker, et al. 2006. “The Epilepsy-Linked Lgi1 Protein Assembles into Presynaptic Kv1 Channels and Inhibits Inactivation by Kvbeta1.” Neuron 49 (5): 697–706. 10.1016/j.neuron.2006.01.033.

Storm, J. F. 1988. “Temporal Integration by a Slowly Inactivating K+ Current in Hippocampal Neurons.” Nature 336 (6197): 379–81. 10.1038/336379a0.

Vaughan, Timothy G. 2017. “IcyTree: Rapid Browser-Based Visualization for Phylogenetic Trees and Networks.” Bioinformatics 33 (15): 2392–94. 10.1093/bioinformatics/btx155.

Waters, Michael F., Natali A. Minassian, Giovanni Stevanin, et al. 2006. “Mutations in Voltage-Gated Potassium Channel KCNC3 Cause Degenerative and Developmental Central Nervous System Phenotypes.” Nature Genetics 38 (4): 447–51. 10.1038/ng1758.

Wei, Aguan, Manuel Covarrubias, Alice Butler, Keith Baker, Michael Pak, and Lawrence Salkoff. 1990. “K+ Current Diversity Is Produced by an Extended Gene Family Conserved in Drosophila and Mouse.” Science 248 (4955): 599–603. 10.1126/science.2333511.

Zhu, Ying, and Fabrizio Gabbiani. 2016. “Fine and Distributed Subcellular Retinotopy of Excitatory Inputs to the Dendritic Tree of a Collision-Detecting Neuron.” Journal of Neurophysiology 115 (6): 3101–12. 10.1152/jn.00044.2016.

